# NAE1-Dependent Protein Neddylation Preserves Endothelial Identity and Vascular Integrity

**DOI:** 10.64898/2026.08.27.746884

**Authors:** Rachelle Nelson Zoni, Fatema Yeasmin Tanni, Chang Min Lee, Xiao Cheng, Jiyao Zhu, Vijaya Bhaskar Baki, MD Sadikul Islam, Jie Li, Huabo Su, Xinghui Sun

**Affiliations:** Department of Biochemistry, University of Nebraska-Lincoln, Beadle Center, 1901 Vine St, Lincoln, Nebraska 68588, USA; Vascular Biology Center, Medical College of Georgia, Augusta University, 1460 Laney Walker Blvd, Augusta, GA 30912; The Nebraska Center for Integrated Biomolecular Communication (NCIBC), University of Nebraska-Lincoln, Lincoln, Nebraska 68588, USA; The Nebraska Center for the Prevention of Obesity Diseases through Dietary Molecules, University of Nebraska-Lincoln, Lincoln, NE 68583, USA

**Author notes:** R.N. Zoni and F.Y. Tanni contributed equally. H. Su and X. Sun contributed equally. Correspondence to: Xinghui Sun, PhD, Department of Biochemistry, University of Nebraska-Lincoln, Beadle Center, 1901 Vine St, Lincoln, Nebraska 68588,; or Huabo Su, PhD, Vascular Biology Center, Medical College of Georgia, Augusta University, 1460 Laney Walker Blvd, Augusta, GA 30912,.

**Keywords:** endothelial cells, protein neddylation, gasdermin, vascular disease, inflammation

## Abstract

**Background:** Endothelial dysfunction is a central driver of cardiovascular and inflammatory diseases, yet the post-translational mechanisms that preserve endothelial homeostasis remain incompletely understood. Protein neddylation, the covalent conjugation of a ubiquitin-like modifier, regulates diverse cellular processes, yet its physiological role in the vascular endothelium remains unknown. This study investigated whether protein neddylation is required to preserve endothelial identity and vascular homeostasis.

**Methods:** We generated tamoxifen-inducible endothelial-specific *Nae1* knockout mice to inhibit neddylation and combined bulk RNA sequencing, single-cell and single-nucleus transcriptomics, quantitative proteomics, biochemical analyses, and gain- and loss-of-function approaches to define the role of endothelial neddylation in vascular homeostasis and inflammatory injury.

**Results:** Endothelial-specific *Nae1* deletion caused rapid mortality associated with vascular leakage, platelet accumulation, inflammation, and multi-organ injury. Multi-omics analyses demonstrated profound loss of endothelial identity, characterized by suppression of core endothelial programs and activation of inflammatory, procoagulant, and pyroptotic pathways. Single-cell analyses revealed progressive endothelial dysfunction culminating in depletion of the endothelial population and remodeling of the vascular niche. Mechanistically, endothelial neddylation deficiency activated gasdermin D (GSDMD)- and gasdermin E (GSDME)-dependent pyroptosis, whereas dual inhibition of GSDMD and GSDME markedly attenuated inflammatory transcriptomic remodeling, vascular injury, hepatocyte death, immune cell infiltration, and platelet accumulation. Translational analyses demonstrated reduced endothelial neddylation in experimental endotoxemia and decreased expression of neddylation pathway components in human atherosclerosis and COVID-19 datasets. Conversely, restoration of endothelial neddylation partially reversed inflammatory endothelial transcriptomic reprogramming in vivo.

**Conclusions:** NAE1-dependent protein neddylation is an essential regulator of endothelial identity and vascular integrity. Loss of endothelial neddylation promotes gasdermin-dependent pyroptosis and thrombo-inflammatory vascular injury, whereas restoration of the neddylation pathway mitigates inflammatory endothelial dysfunction. These findings identify endothelial neddylation as a fundamental mechanism maintaining vascular homeostasis and a potential therapeutic target for cardiovascular and inflammatory diseases.

**Clinical Perspective:** What Is New?

- Endothelial-specific deletion of NAE1, the enzyme that initiates protein neddylation, causes rapid mortality in mice from multi-organ tissue damage, cell death, and inflammation.
- Neddylation loss activates the pyroptosis executioners GSDMD and GSDME, and silencing GSDMD and GSDME together reverses the transcriptomic and tissue-level damage caused by neddylation deficiency.
- The endothelial neddylation pathway is significantly downregulated in human clinical conditions, including atherosclerosis and COVID-19, as well as in experimental models of systemic inflammation.

What Are the Clinical Implications?

- Reduced neddylation in atherosclerotic human arteries suggests this pathway could serve as a biomarker or intervention point for endothelial dysfunction in cardiovascular disease.
- GSDMD and GSDME represent potential therapeutic targets for protecting endothelial integrity in vascular and inflammatory diseases, including endotoxemia.
- Because neddylation inhibitors such as MLN4924 (pevonedistat) are already in clinical use for cancer, these findings suggest a mechanism behind their reported vascular and hepatic toxicity that warrants monitoring.

## Introduction

Vascular endothelial cells (ECs) are indispensable for maintaining tissue homeostasis by preserving vascular barrier integrity, regulating leukocyte trafficking, preventing thrombosis, and coordinating inflammatory responses^1,2^. Endothelial function is tightly controlled through coordinated epigenetic, transcriptional, and post-translational mechanisms that enable rapid adaptation to environmental cues^3–8^. During inflammation, infection, or metabolic stress, ECs undergo activation characterized by increased expression of adhesion molecules, including E-selectin and vascular cell adhesion molecule-1 (VCAM-1)^9^, which promote leukocyte recruitment to sites of injury. Concurrent disruption of endothelial junctions, particularly through loss of VE-cadherin, compromises vascular barrier integrity and increases permeability^10,11^. Endothelial injury also promotes platelet activation and coagulation, contributing to thrombo-inflammatory responses^12^. Although these responses are initially adaptive, persistent endothelial dysfunction drives the development of obesity, insulin resistance, diabetes, metabolic dysfunction-associated steatotic liver disease, hypertension, atherosclerosis, and other cardiovascular disorders^2,13,14^. Defining the molecular mechanisms that preserve endothelial homeostasis therefore remains a major goal for developing therapies for vascular disease.

Protein post-translational modifications provide a rapid and energy-efficient mechanism for regulating protein abundance and function beyond transcriptional control. Indeed, transcriptomic and proteomic studies have demonstrated only modest concordance between mRNA and protein abundance across human tissues^15,16^, underscoring the importance of post-translational regulation in cellular physiology. Among these modifications, ubiquitination^17^ and SUMOylation^18,19^ are established regulators of endothelial biology, whereas the contribution of other ubiquitin-like modifiers remains poorly understood^20,21^. NEDD8 (neural precursor cell expressed, developmentally downregulated 8) is an evolutionarily conserved ubiquitin-like protein that modifies target proteins through the reversible process of neddylation^22,23^. Neddylation is initiated by the NEDD8-activating enzyme, composed of the catalytic subunit UBA3 and the regulatory subunit NAE1, and proceeds through a dedicated E1–E2–E3 enzymatic cascade to conjugate NEDD8 to lysine residues on substrate proteins^22,23^, most notably Cullin family proteins, thereby activating Cullin-RING E3 ubiquitin ligases (CRLs). This modification regulates protein stability, localization, and activity and thereby controls numerous cellular processes^22–25^. Genetic and pharmacological studies have established critical roles for neddylation in neurodegeneration^26,27^, immune disorders^28^, and cardiovascular development^29–32^, and pharmacological inhibition of neddylation has demonstrated potent therapeutic effects in malignancies^33^. Nevertheless, whether NAE1-dependent neddylation is required to preserve endothelial homeostasis and vascular integrity remains unknown.

Maintenance of endothelial identity and survival is essential for vascular homeostasis. Loss of endothelial identity has been reported in multiple pathological conditions, including sepsis and cardiovascular disease^34–36^, and is associated with impaired vascular function. Although EC death has traditionally been attributed to apoptosis, accumulating evidence indicates that pyroptosis, an inflammatory form of programmed cell death mediated by gasdermin (GSDM) proteins, also contributes to vascular injury^37,38^. Pyroptosis is initiated by inflammatory signaling pathways that activate caspases and induce proteolytic cleavage of GSDMs^39,40^, leading to membrane pore formation, cell lysis, and amplification of inflammation^41,42^. While the molecular regulation of pyroptosis has been extensively studied in immune cells, the mechanisms governing endothelial pyroptosis remain incompletely understood^43^.

In this study, we demonstrate that endothelial neddylation is indispensable for vascular integrity and postnatal survival. Utilizing a tamoxifen-inducible, endothelial-specific *Nae1* knockout mouse model, we show that the loss of neddylation leads to rapid mortality, accompanied by cell death, vascular leakage, inflammation, and multi-organ injury. Integrative bulk RNA sequencing, single-cell and single-nucleus transcriptomics, quantitative proteomics, and biochemical analyses reveal that neddylation-deficient ECs undergo a profound loss of endothelial identity, characterized by the downregulation of core vascular markers and the activation of inflammatory, pro-coagulant, and pyroptotic transcriptional programs; and identify GSDM-dependent pyroptosis as a major downstream consequence of impaired endothelial neddylation. Genetic inhibition of GSDMD and GSDME markedly attenuates transcriptomic remodeling, vascular injury, and tissue inflammation. Finally, we show that endothelial neddylation is suppressed in experimental endotoxemia and human vascular disease, and restoration of the neddylation pathway partially reverses inflammatory endothelial reprogramming. Together, these findings identify NAE1-mediated protein neddylation as an essential regulator of endothelial homeostasis, establish impaired neddylation as a previously unrecognized mechanism driving vascular dysfunction and inflammatory tissue injury, and suggest that restoring this pathway may offer a novel therapeutic strategy for treating inflammatory and cardiovascular diseases.

## Methods

### Data Availability

The detailed methods and materials and the Major Resources Table are provided in the Supplemental Material. All data generated or analyzed in this study are included in this published article and its supplementary information files. RNA-sequencing data from this study are available through the Gene Expression Omnibus under accession numbers GSE336753, GSE336749, GSE336746, and GSE336754.

### Experimental animals

All studies were approved by the Institutional Animal Care and Use Committee at the University of Nebraska–Lincoln. *Nae1*^fl/fl^ mice ^29^ were crossed with *Cdh5-CreERT2* mice ^44,45^ to generate *Nae1*^fl/fl^; Cdh5^CreERT2+^ mice (referred to as *Nae1* iECKO mice), in which *Nae1* was specifically deleted in the vascular endothelium after tamoxifen treatment. Tamoxifen (Sigma T5648) was administered intraperitoneally (50 mg/kg/day) into *Nae1*^fl/fl^ and *Nae1* iECKO mice at four – six weeks of age for five consecutive days. Both male and female mice were used for experiments, and tissues were collected at early (14 days) or late time points (25-28 days) post-tamoxifen injection for downstream analysis. Blood was collected through cardiac puncture and supplemented with 4 mM EDTA or 3.2% buffered sodium citrate (1:9 volume ratio) to obtain plasma. After perfusion, the same anatomic lobe of the liver or lung from each mouse was fixed with 10% neutral buffered formalin at 4°C for 24 hours, and other lobes were snap-frozen for storage at -80°C. Fixed liver tissues were processed for OCT embedding and cryo-sectioning or paraffin embedding and sectioning. Perfusion fixation of lungs was performed in some mice for histological analysis. Other tissues were appropriately collected and stored.

### Statistical Analysis

Normality was assessed using the Shapiro-Wilk test, and variance homogeneity was assessed using the Levene test. Two-group comparisons were performed using Student’s *t* test or Welch’s *t* test for normally distributed data, and the Mann-Whitney *U* test for data not normally distributed. For multigroup comparisons, a two-way ANOVA with Tukey’s HSD post-hoc test was applied for experiments involving two variables. Sample sizes denote biological replicates unless otherwise stated. Differences in survival were assessed using the Kaplan-Meier log-rank test. Data are presented as mean ± SEM. Statistical significance was defined as *P*<0.05. Analyses were performed using R 4.4.3.

## Results

### Loss of endothelial neddylation triggers tissue injury, inflammation, and mortality in mice

To define the physiological role of endothelial neddylation, we generated tamoxifen-inducible endothelial-specific *Nae1* knockout mice (Nae1^fl/fl^; *Cdh5^CreERT^*^2+^, hereafter referred to as *Nae1* iECKO) (Figure 1A). Tamoxifen administration efficiently deleted *Nae1*, which encodes the regulatory subunit of the NEDD8-activating E1 enzyme, in hepatic endothelial cells (ECs). This reduced NAE1 protein expression by 51.1% and decreased total neddylated cullins and neddylated Cullin-3 by 74.1% and 79.3%, respectively, confirming efficient inhibition of neddylation (Figures 1B and S1A). Endothelial loss of neddylation caused rapid mortality, with all *Nae1* iECKO mice dying between 30 and 42 days after tamoxifen administration (Figure 1C), despite no change in body weight (Figure S1B).

**Figure 1.**
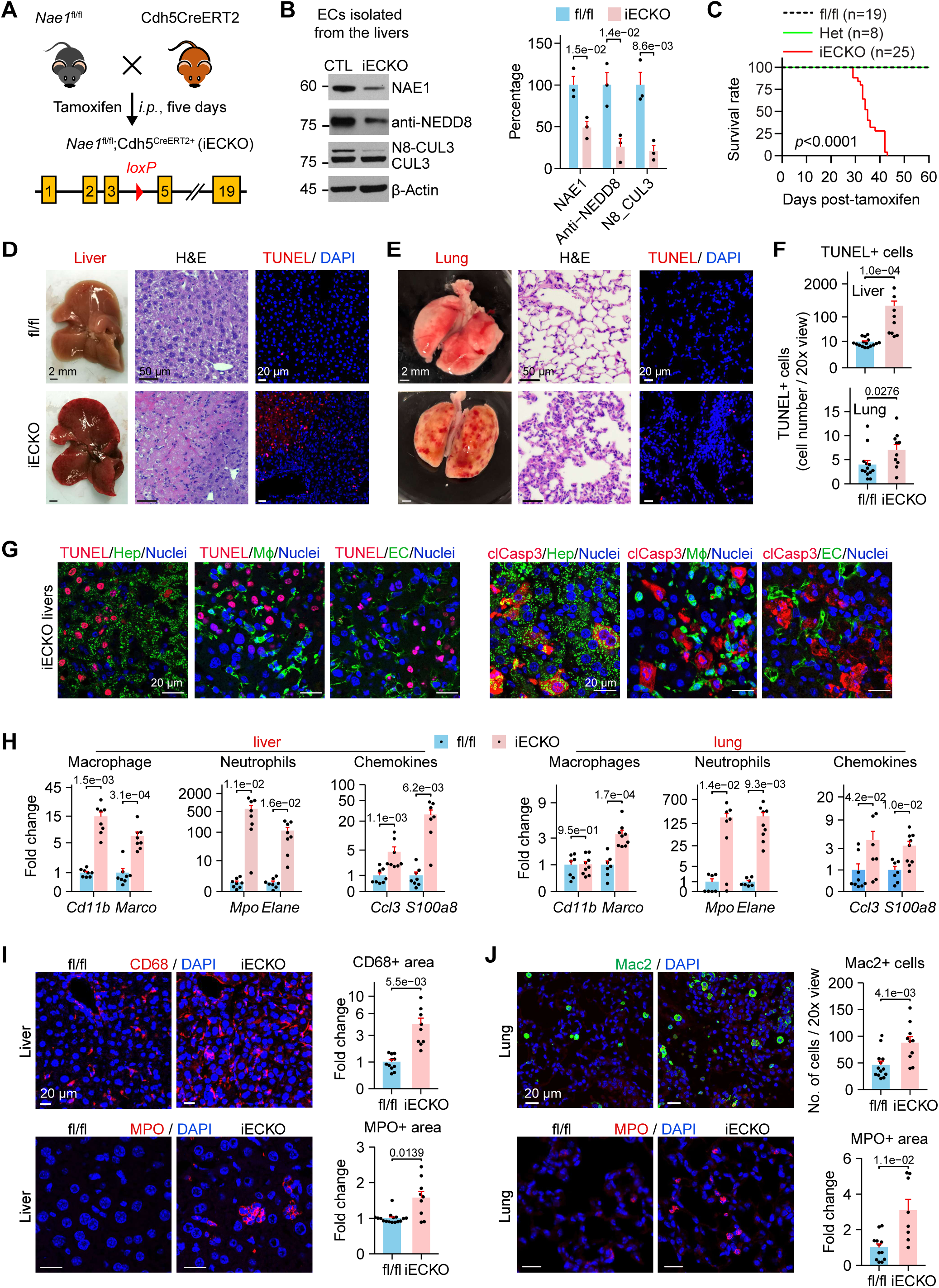
Loss of endothelial neddylation triggers tissue injury, inflammation, and mortality in mice. **A**, Schematic depicting the generation of tamoxifen-inducible EC-specific *Nae1* knockout mice (*Nae1* iECKO). Cdh5^CreERT2^ mice express Cre driven by the promoter of VE-cadherin (Cdh5) gene upon tamoxifen administration. **B**, Western blot for NAE1, NEDD8, and CUL3 in liver ECs isolated from *Nae1* fl/fl and iECKO mice 28 days after tamoxifen administration. **C**, Survival curve after tamoxifen administration. **D**, Gross anatomy images, Hematoxylin and Eosin (H&E) staining, and TUNEL staining showing the livers and liver sections of *Nae1* fl/fl and iECKO mice. **E**, Gross anatomy images, Hematoxylin and Eosin (H&E) staining, and TUNEL staining showing the lungs and lung sections of *Nae1* fl/fl and iECKO mice. **F**, Quantification of TUNEL staining in the livers and lungs of *Nae1* fl/fl and iECKO mice. **G**, TUNEL and immunofluorescence double-labeling assay for detecting apoptotic cells. **H**, The levels of mRNA expression of indicated genes by quantitative RT-qPCR in the liver and lungs of *Nae1* fl/fl and iECKO mice. **I-J**, Immunofluorescence staining of macrophage marker proteins (CD68 and Mac2), and neutrophil marker protein (MPO) in the livers and lungs of fl/fl or iECKO mice. Statistical significance (*P* < 5.0e-2) was determined by the Student’s *t-*test (B, lung *Cd11b* in H), the Mann-Whitney *U* test (F; liver *Marco* and *Ccl3* in H; lung *Marco*, *Mpo*, and *Ccl3* in H; liver MPO in I; and Mac2 in J), the Welch’s *t*-test (liver *Cd11b*, *Mpo*, *Elane*, and *S100a8* in H; lung *Elane* and *S100a8* in H; CD68 in I, and lung MPO in J), and Kaplan-Meier log-rank test (C).

Gross examination and hematoxylin and eosin staining revealed extensive hemorrhage, necrosis, and inflammatory lesions in the livers and lungs of *Nae1* iECKO mice (Figure 1D-E). TUNEL staining demonstrated a 28-fold increase in TUNEL-positive cells in the liver and a 1.8-fold increase in the lung compared with control mice (Figure 1D-F). Dual immunofluorescence of livers showed that TUNEL-positive cells were predominantly hepatocytes, whereas ECs and macrophages were largely TUNEL-negative, indicating that endothelial *Nae1* deficiency primarily induces secondary hepatocyte death. Cleaved caspase-3 immunostaining further confirmed extensive hepatocyte apoptosis (Figure 1G).

Because tissue injury is typically accompanied by inflammatory cell recruitment, we next examined immune cell infiltration. Endothelial *Nae1* deletion markedly increased the expression of macrophage- and neutrophil-associated marker genes, together with multiple chemokines, in both the liver and lung (Figure 1H). Immunostaining confirmed increased accumulation of macrophages (CD68 and galectin-3/Mac2) and neutrophils (myeloperoxidase [MPO]) in both organs (Figures 1I and 1J). Similarly, immune cell marker expression was increased in the aorta but not in the heart (Figures S1D and S1E), indicating organ-specific vascular inflammation. Cardiac function remained unchanged (Figure S1F).

These findings establish endothelial neddylation as an essential regulator of postnatal survival and vascular homeostasis. Endothelial *Nae1* deletion causes fatal liver and lung injury characterized by extensive cell death, inflammatory cell infiltration, and vascular inflammation.

### Endothelial neddylation deficiency reprograms the endothelial transcriptome and proteome

To define the early molecular consequences of endothelial neddylation deficiency, we performed bulk RNA sequencing of CD31+ and CD146+ hepatic ECs 14 days after tamoxifen administration (Figure 2A). Comparison of independently isolated EC populations identified 618 shared differentially expressed genes (DEGs) between *Nae1* fl/fl and *Nae1* iECKO mice (Figures 2B and S2A). Gene set enrichment analysis demonstrated significant enrichment of the vascular endothelial core gene set (GSE210119)^7^ in *Nae1* fl/fl ECs, accompanied by marked downregulation of key endothelial genes, including *Kdr* and *Flt1* (Figures 2C and S2B). Ingenuity Pathway Analysis further revealed enrichment of pathways associated with phagosome formation, neutrophil extracellular trap signaling, leukocyte extravasation, TREM1 signaling, natural killer cell signaling, NRF2-mediated oxidative stress responses, pyroptosis, HIF-1α signaling, thrombin signaling, and growth factor-regulated epithelial-to-mesenchymal transition (Figure 2D). Transcription factor activity analysis also predicted suppression of the endothelial identity regulator ERG^7^ together with activation of inflammatory transcription factors, including SPI1^46^ and CEBPD^47^ (Figure S2C). Together, these findings indicate that neddylation-deficient ECs lose endothelial identity and acquire inflammatory, stress-responsive, procoagulant, and mesenchymal-like transcriptional programs.

**Figure 2.**
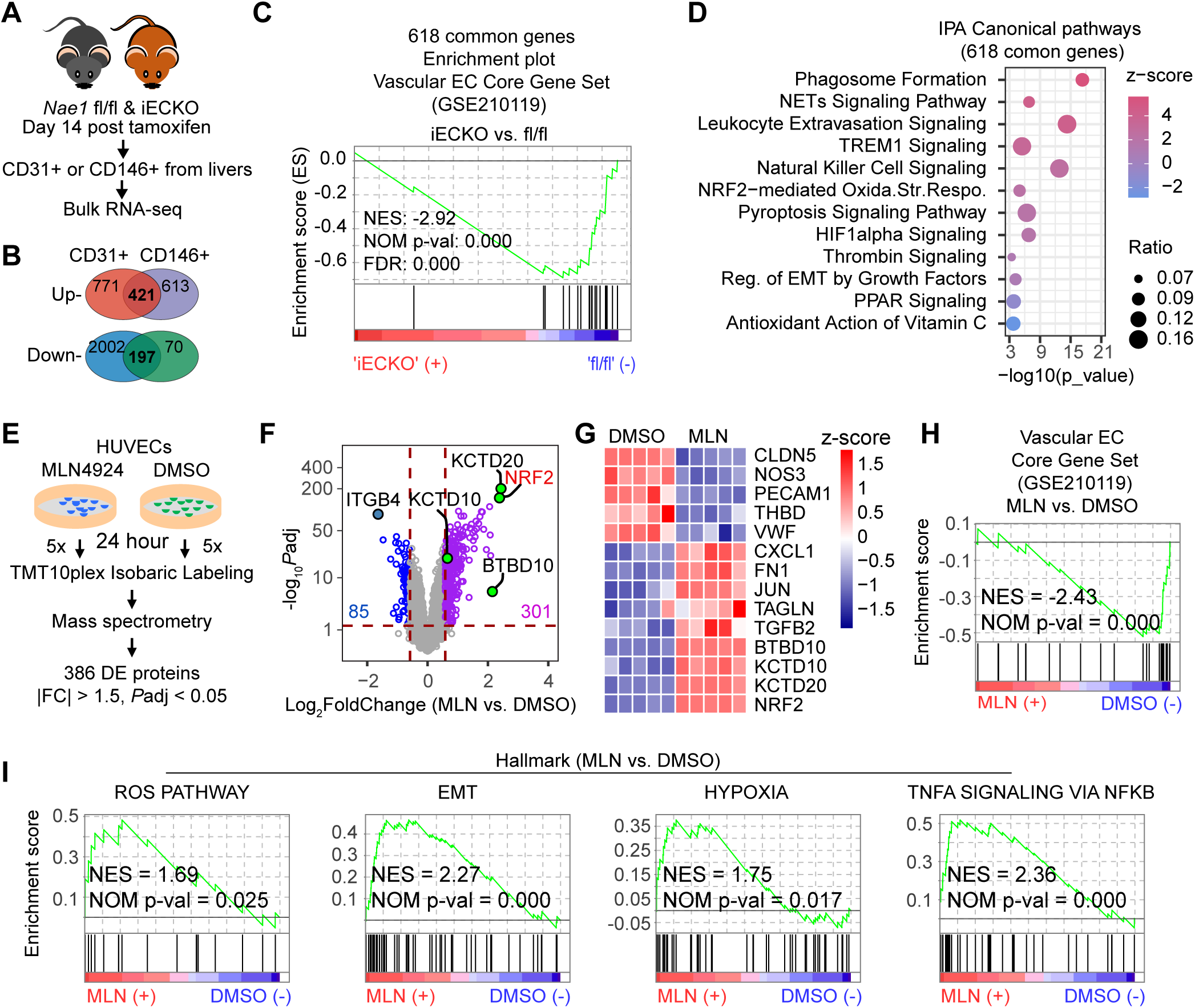
Endothelial neddylation deficiency reprograms the endothelial transcriptome and proteome. **A**, Schematic depicting the workflow of bulk RNA-seq analysis. CD31+ or CD146+ cells were isolated from 14 livers (CD31+; n=6 for the fl/fl samples, and n=8 for the iECKO samples) or 16 livers (CD146+; n=8 for the fl/fl and iECKO samples, respectively) of mice 14 days after tamoxifen administration (50mg/kg, five consecutive daily *i.p.*). **B**, A Vienn Diagram shows the common 618 DEGs (*P*adj <0.05; |log2FoldChange|>0.585) identified by bulk RNA-seq of CD31+ and CD146+ cells. **C**, The vascular EC Core Gene Set^7^ (GSE210119) was enriched in the fl/fl phenotype. The common 618 DEGs were used for GSEA. Normalized counts from CD31+ bulk RNA-seq were used. **D**, The common 618 DEGs were used for Ingenuity Pathway Analysis (IPA). Circle size reflects the ratio that measures the proportion of input genes mapped to each individual pathway term, whereas color represents the z-score that predicts the activation (plus z-score) or inhibition (minus z-score) of the pathway. The x-axis represents the minus log10-transformed p-values. **E**, Schematic depicting the workflow of quantitative proteomics by TMT10 plex labeling followed by mass spectrometry and analysis. HUVECs were treated with DMSO or MLN4924 (MLN) at 100 nM for 24 hours (n=5 biological replicates / condition). **F**, A volcano plot showing the differentially expressed protein identified by the quantitative proteomics (|FC| > 1.5 and *P*adj < 0.05). **G**, A heatmap showing the relative expression of EC identity proteins (gene symbols are CLDN5, NOS3, PECAM1, THBD, and VWF), inflammatory mediators (CXCL1, FN1, JUN, TAGLN), and Cullin-3 adaptor/target proteins (BTBD10, KCTD10, KCTD20, and NRF2). **H and I**, The vascular EC Core Gene Set^7^ (GSE210119) enriched in DMSO-treated HUVECs and other Hallmark gene sets enriched in MLN4924-treated HUVECs. The UniProt ID mapping tool was used to convert protein IDs to gene symbols. Normalized protein expression was used for GSEA.

To define the global proteomic consequences of neddylation inhibition, human umbilical vein endothelial cells (HUVECs) were treated with the NEDD8-activating enzyme inhibitor MLN4924 (100 nM) or vehicle for 24 hours (Figure 2E). Tandem mass tag (TMT)-based quantitative proteomics identified 301 upregulated and 85 downregulated proteins following MLN4924 treatment (Figure 2F). Consistent with inhibition of Cullin-RING ligase activity, NRF2, a canonical Cullin-3 substrate, was among the most strongly increased proteins. In contrast, several proteins critical for endothelial function, including CLDN5, NOS3, PECAM1, THBD, and VWF, were reduced (Figure 2G), with many belonging to the vascular endothelial core gene set (Figure 2H). Conversely, proteins associated with inflammation and endothelial-to-mesenchymal transition, including CXCL1, FN1, JUN, and TAGLN, were increased.

Multiple Cullin-3 adaptors or substrates, including NRF2^48^, BTBD10^49^, KCTD10^49^, and KCTD20^49^, also accumulated, consistent with impaired Cullin-3 activity. Gene set enrichment analysis further demonstrated enrichment of hallmark pathways related to oxidative stress, hypoxia, epithelial-to-mesenchymal transition, and inflammation (Figure 2I).

Collectively, transcriptomic and proteomic analyses demonstrate that NAE1-dependent neddylation is required to preserve endothelial homeostasis by maintaining EC identity while restraining inflammatory, stress-response, and mesenchymal phenotypic reprogramming.

### Endothelial neddylation deficiency activates endothelial cells, disrupts vascular integrity, and promotes platelet aggregation

Because endothelial activation is a hallmark of vascular injury^1^, we next examined the effects of *Nae1* deletion on endothelial activation and barrier function. In the liver, endothelial *Nae1* deletion increased both mRNA and protein expression of the adhesion molecules VCAM-1 and E-selectin (*Sele*), whereas in the lung only E-selectin was significantly increased (Figures 3A–3C), indicating tissue-specific endothelial activation.

**Figure 3.**
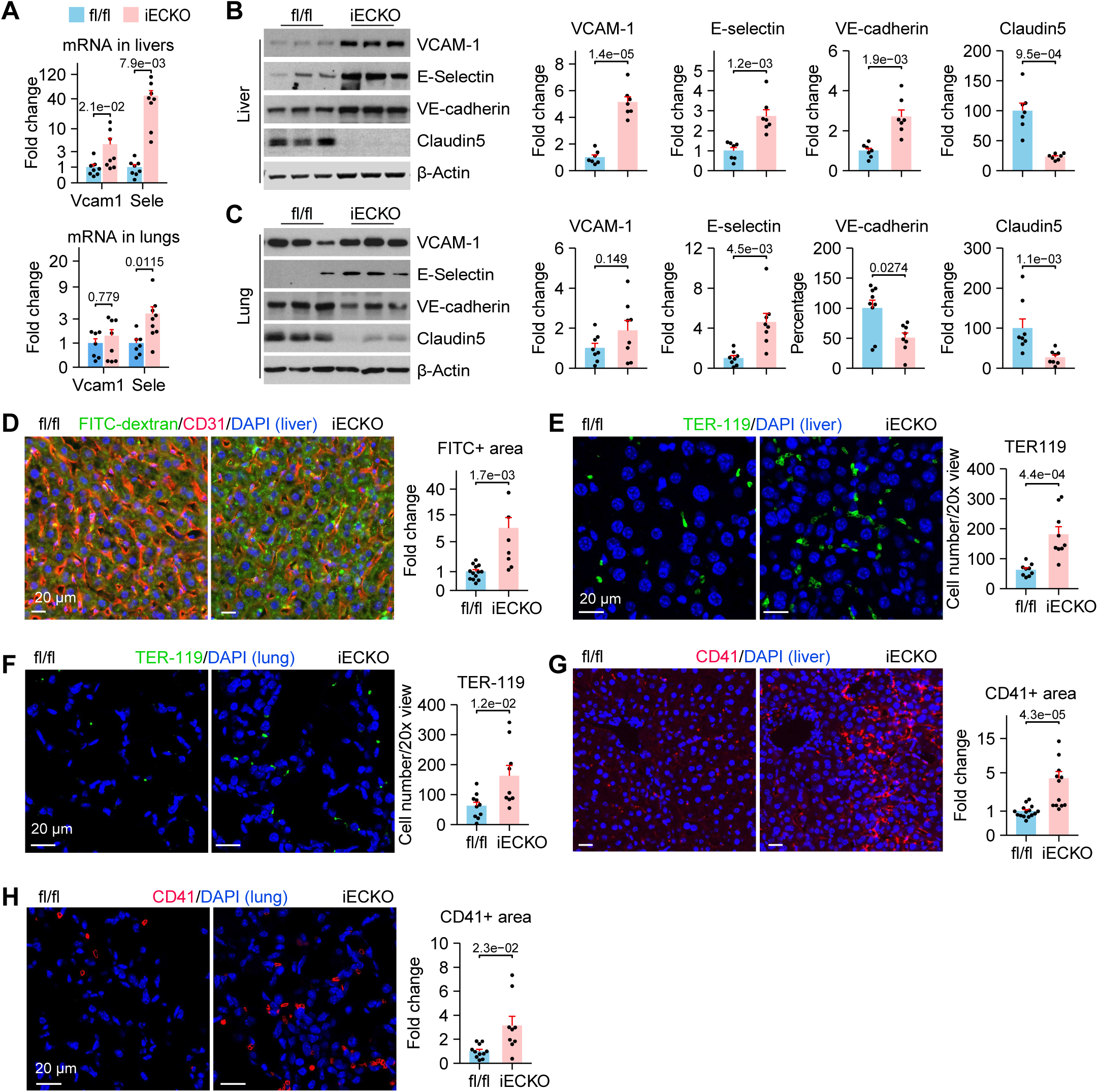
Endothelial neddylation deficiency activates endothelial cells, disrupts vascular integrity, and promotes platelet aggregation. **A**, *Vcam1* and E-selectin (*Sele*) mRNA expression assessed by qPCR in the lungs and livers of *Nae1* fl/fl and iECKO mice at day 28 post-tamoxifen injection. **B and C**, VCAM-1, E-selectin, VE-cadherin, Claudin-5 protein expression assessed by Western Blot and quantified in the livers (B) and lungs (C) of *Nae1* fl/fl and iECKO mice at day 28 post-tamoxifen injection. **D**, FITC-dextran (70 kDa) extravasation was measured to reflect vascular permeability. FITC-dextran was intravenously administered (10 mg/mouse), and tissues were collected at 30 min after injection. **E and F**, Immunofluorescence staining and quantification of erythroid cell marker protein (TER-119) in the livers (E) and lungs (F) of fl/fl or iECKO mice. **G and H**, Immunofluorescence staining and quantification of platelet marker protein (CD41) in the livers (G) and lungs (H) of fl/fl or iECKO mice. Statistical significance (*P* < 5.0e-2) was determined by the Student’s *t-*test (VCAM-1, E-selectin, VE-cadherin in B; lung E-selectin in C; E, and F), the Welch’s *t*-test (liver *Sele* in A, liver Claudin5 in B, lung VCAM-1 in C, and CD41 in H), and the Mann-Whitney *U* test (lung *Vcam1* and *Sele* and liver *Vcam1* in A; lung VE-cadherin and Claudin5 in C; D, and G).

We next assessed endothelial junctional proteins that maintain vascular barrier integrity. Claudin-5 protein expression was markedly reduced in both the liver and lung of *Nae1* iECKO mice (Figures 3B and 3C). VE-cadherin expression decreased in the lung but, unexpectedly, increased in the liver. Despite this difference, FITC-dextran extravasation demonstrated significantly increased vascular permeability in the liver following endothelial *Nae1* deletion (Figure 3D), indicating impaired endothelial barrier function.

Consistent with vascular leakage, TER-119^50^ immunostaining revealed increased erythrocyte extravasation in both the liver and lung of *Nae1* iECKO mice (Figures 3E and 3F), supporting the presence of leaky vessels and hemorrhage^51^. CD41 immunostaining further demonstrated prominent platelet aggregation in both organs (Figures 3G and 3H), indicative of a prothrombotic vascular phenotype^52^.

Together, these findings demonstrate that endothelial neddylation is required to maintain endothelial quiescence and vascular barrier integrity. Loss of *Nae1* promotes endothelial activation, vascular leakage, hemorrhage, and platelet aggregation, establishing a proinflammatory and procoagulant vascular microenvironment.

### Single-cell transcriptomic analyses reveal progressive loss of endothelial identity following endothelial *Nae1* deletion

To define the temporal and molecular consequences of endothelial neddylation deficiency at single-cell resolution, we first performed single-cell RNA sequencing (scRNA-seq) of CD31+ cells isolated from mouse livers 14 days after tamoxifen administration (Figure 4A). Unsupervised clustering identified five major cell populations (Figures 4B and S3B). Although the overall cellular composition was largely preserved, *Nae1* deletion increased the proportion of macrophages from 10.9% to 17.4% while reducing the EC population from 69.3% to 63.4%. Consistent with enrichment of the NRF2 signaling pathway and increased NRF2 expression in neddylation-deficient ECs (Figure 2D, G), several canonical NRF2 target genes, including *Gclm*, *Nqo1*, *Gclc*, and *Gsr*, were among the most highly upregulated transcripts, indicating robust NRF2 activation resulting from marked disruption of Cullin-3 activity (Figure 4C). Expression of Ninj1, a mediator of pyroptotic cell death^53^, was also increased. Gene set enrichment analysis further demonstrated enrichment of the vascular endothelial core gene set in control ECs, whereas genes associated with pyroptosis were preferentially enriched in *Nae1* iECKO ECs (Figure S3A), indicating early loss of endothelial identity accompanied by activation of inflammatory cell death programs.

**Figure 4.**
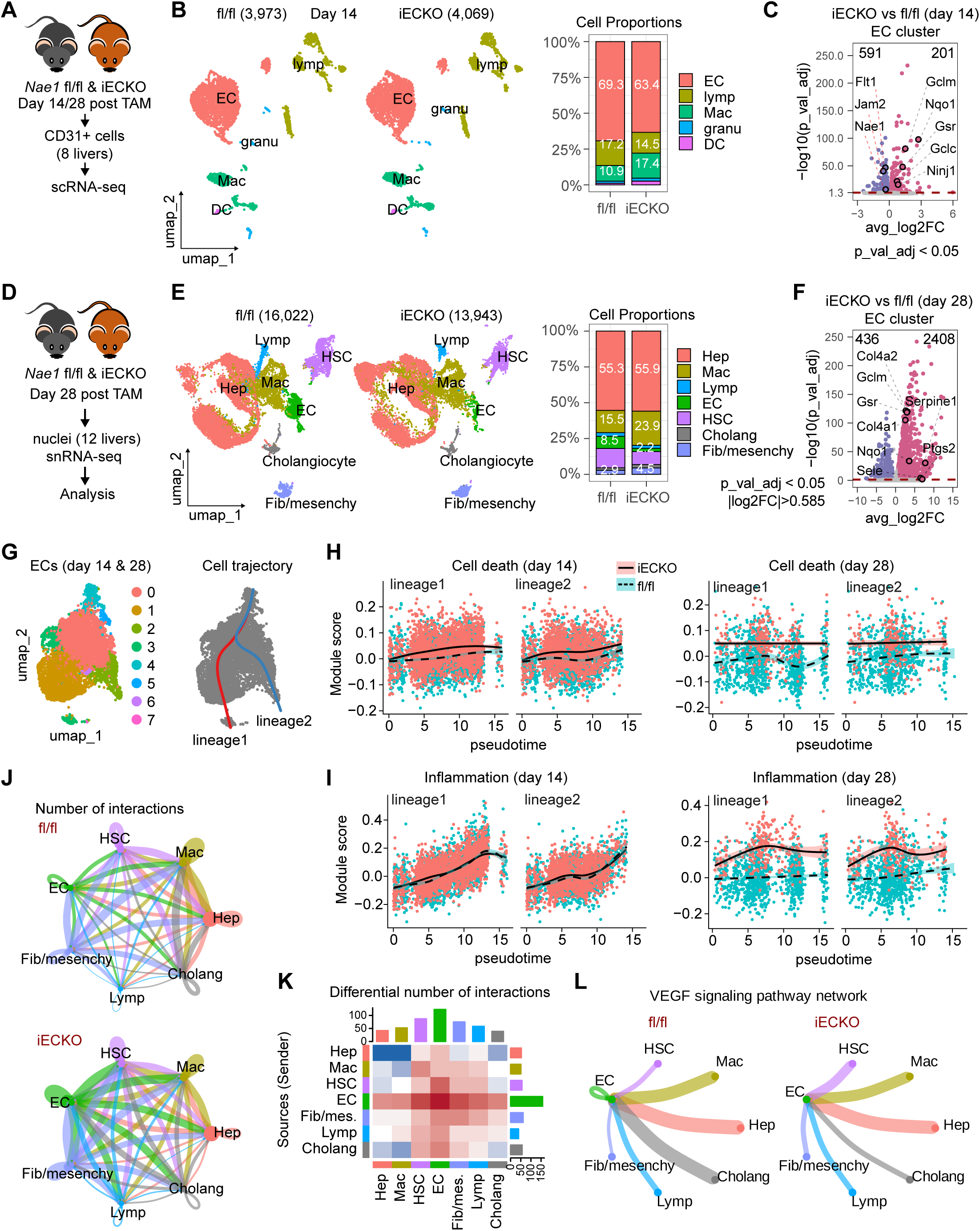
Single-cell transcriptomic analyses reveal progressive loss of endothelial identity following endothelial *Nae1* deletion. **A**, Schematic depicting the workflow of single-cell RNA-seq (scRNA-seq) analysis. CD31+ cells were isolated from 8 livers (n=4 for the fl/fl and iECKO samples, respectively) of mice at day 14 (or day 28) after tamoxifen injection (50mg/kg, five consecutive daily *i.p.*). **B**, Uniform manifold approximation and projection (UMAP) of scRNA-seq for the fl/fl (n=3,973 cells) and iECKO (n=4,069 cells) samples after quantity control and data filtering using Harmony integration. Cell proportion of annotated cell clusters in the fl/fl and iECKO samples were shown. **C**, Differentially expressed genes (DEGs; |avg_log2FC| > 0.585 and p_val_adj < 0.05) identified by the FindMarkers function in the Seurat package using the Wilcoxon Rank Sum test. **D**, Schematic depicting the workflow of single-nuclei RNA-seq (snRNA-seq) analysis. Nuclei were isolated from 12 livers (n =6 for the fl/fl and iECKO samples, respectively) of mice at day 28 after tamoxifen administration (50mg/kg, five consecutive daily *i.p.*). **E**, Uniform manifold approximation and projection (UMAP) of snRNA-seq for the fl/fl (n=16,022 nuclei) and iECKO (n=13,943 nuclei) samples after quantity control and data filtering using Harmony integration. Cell proportion of annotated cell clusters in the fl/fl and iECKO samples were shown. **F**, Differentially expressed genes (DEGs; |avg_log2FC| > 0.585 and p_val_adj < 0.05) identified by the FindMarkers function in the Seurat package using the Wilcoxon Rank Sum test. **G**, Uniform manifold approximation and projection (UMAP) of EC populations from both scRNA-seq and snRNA-seq (left) and cell lineages revealed by cell trajectory analysis using Slingshot (right). **H and I**, The cell death and inflammation module scores were plotted as the function of pseudotime from cell trajectory analysis. The module scores were calculated using the AddModuleScore function in Seurat, which measure the average expression levels of genes related to cell death (H) and inflammation (I). **J**, The numbers of biologically significant cell-cell communication networks inferred by CellChat. **K**, A heatmap displays the differential number of cell-cell interactions from CellChat analysis among the indicated populations. The top-colored bar plot represents the sum of column of values displayed in the heatmap (incoming signaling). The right-colored bar plot represents the sum of row of values (outgoing signaling). Red (blue) color in the heatmap represents increased (or decreased) signaling in the iECKO sample compared to the fl/fl sample. **L**, The circle plots show both autocrine and paracrine interactions of VEGF signaling pathway network. A line looping back into the same cell group shows autocrine signaling (e.g., the green line in ECs). Line colors match the sending (source) cell group. For example, the green line indicates that the signaling is from ECs. The line width represents the strength of interactions between two cell groups.

By day 28 after tamoxifen administration, the EC cluster was no longer detected by scRNA-seq in *Nae1* iECKO livers (Figure S3B and S3C), suggesting extensive endothelial transcriptional remodeling and/or depletion. To characterize these later-stage changes before the onset of mortality, we performed single-nucleus RNA sequencing (snRNA-seq) on liver samples collected on day 28 (Figure 4D). Analysis of 16,022 nuclei from *Nae1* fl/fl livers and 13,943 nuclei from *Nae1* iECKO livers identified seven major cell populations, including hepatocytes, macrophages, hepatic stellate cells, ECs, fibroblast/mesenchymal cells, lymphocytes, and cholangiocytes (Figures 4E and S3D). Compared with controls, the proportion of ECs decreased markedly from 8.5% to 2.2%, whereas macrophages and fibroblast/mesenchymal cells increased from 15.5% to 23.9% and from 2.9% to 4.5%, respectively (Figures 4E and S3D), consistent with progressive endothelial loss and tissue remodeling.

Differential expression analysis identified 2,844 differentially expressed genes within the EC population (Figure 4F). As observed at day 14, NRF2 target genes remained among the most strongly induced transcripts, confirming sustained disruption of Cullin-3–dependent ubiquitin ligase activity. Basement membrane genes (*Col4a1* and *Col4a2*) and inflammatory genes, including *Serpine1*, *Sele*, and *Ptgs2*, were also significantly upregulated (Figure 4F). Gene set enrichment analysis demonstrated enrichment of hallmark pathways related to mitotic spindle, apical junction, epithelial-to-mesenchymal transition, reactive oxygen species, hypoxia, and TNF-α signaling via NF-κB in iECKO ECs, whereas pathways involved in oxidative phosphorylation, fatty acid metabolism, xenobiotic metabolism, and other metabolic processes were preferentially enriched in control ECs (Figure S3E).

Together, these single-cell transcriptomic analyses demonstrate that endothelial *Nae1* deletion initiates a progressive program of endothelial dysfunction characterized by early loss of endothelial identity, activation of NRF2-dependent stress responses and pyroptotic signaling, and subsequent depletion of the endothelial population accompanied by inflammatory and mesenchymal remodeling.

To further define the dynamic effects of endothelial neddylation deficiency, we integrated the EC populations (6,996 cells) identified by day-14 scRNA-seq and day-28 snRNA-seq and performed clustering, trajectory inference, pathway activity analysis, and cell-cell communication analysis (Figures 4G–4L, S3F, and S4). Integrated analysis identified two major endothelial lineages (Figure 4G), enabling reconstruction of the progressive phenotypic changes induced by *Nae1* deletion.

At day 14, *Nae1* iECKO ECs exhibited a modest increase in cell death module scores, whereas inflammatory module scores remained comparable to those of control ECs along both endothelial lineages (Figures 4H and 4I). By day 28, however, both cell death and inflammatory signatures were markedly increased throughout the endothelial population, indicating progressive activation of inflammatory cell death programs during disease progression.

To determine how endothelial dysfunction reshapes the hepatic microenvironment, we analyzed intercellular communication using CellChat. Compared with control livers, *Nae1* iECKO livers exhibited substantially increased signaling between ECs and neighboring hepatic cell populations (Figures 4J, 4K, and S4A), consistent with extensive remodeling of the vascular niche. Pathway-specific analysis revealed marked attenuation of VEGF signaling (ECs ◊ ECs and cholangiocytes ◊ ECs), a pathway essential for endothelial homeostasis, together with enhanced signaling through extracellular matrix remodeling pathways, including fibronectin (FN1), collagen, laminin, and bone morphogenetic protein (BMP) pathways (Figures 4L, S4B, and S4C).

These integrated analyses demonstrate that endothelial neddylation deficiency drives a progressive transition from early endothelial stress to widespread inflammatory remodeling. As disease progresses, ECs acquire inflammatory cell death programs, exhibit impaired VEGF-mediated homeostatic signaling, and actively remodel their extracellular microenvironment through enhanced matrix-associated signaling, ultimately contributing to loss of vascular integrity.

### Endothelial neddylation deficiency activates gasdermin-mediated pyroptosis in the hepatic endothelium

GSDMs are the executioners of pyroptosis^54,55^. Because transcriptomic analyses consistently implicated pyroptosis following endothelial *Nae1* deletion (Figures 2D, 4H, and S3A), we examined activation of the GSDM pathway in the liver. Immunoblot analysis of whole-liver lysates demonstrated increased expression of GSDMD, GSDME, and the inflammasome component NLRP3^56^ (NACHT-, LRR- and pyrin domain-containing protein 3) in *Nae1* iECKO mice (Figure 5A and 5B). However, cleavage of GSDMD and GSDME was not affected by *Nae1* deletion, suggesting that pyroptosis occurs in a restricted cellular population.

**Figure 5.**
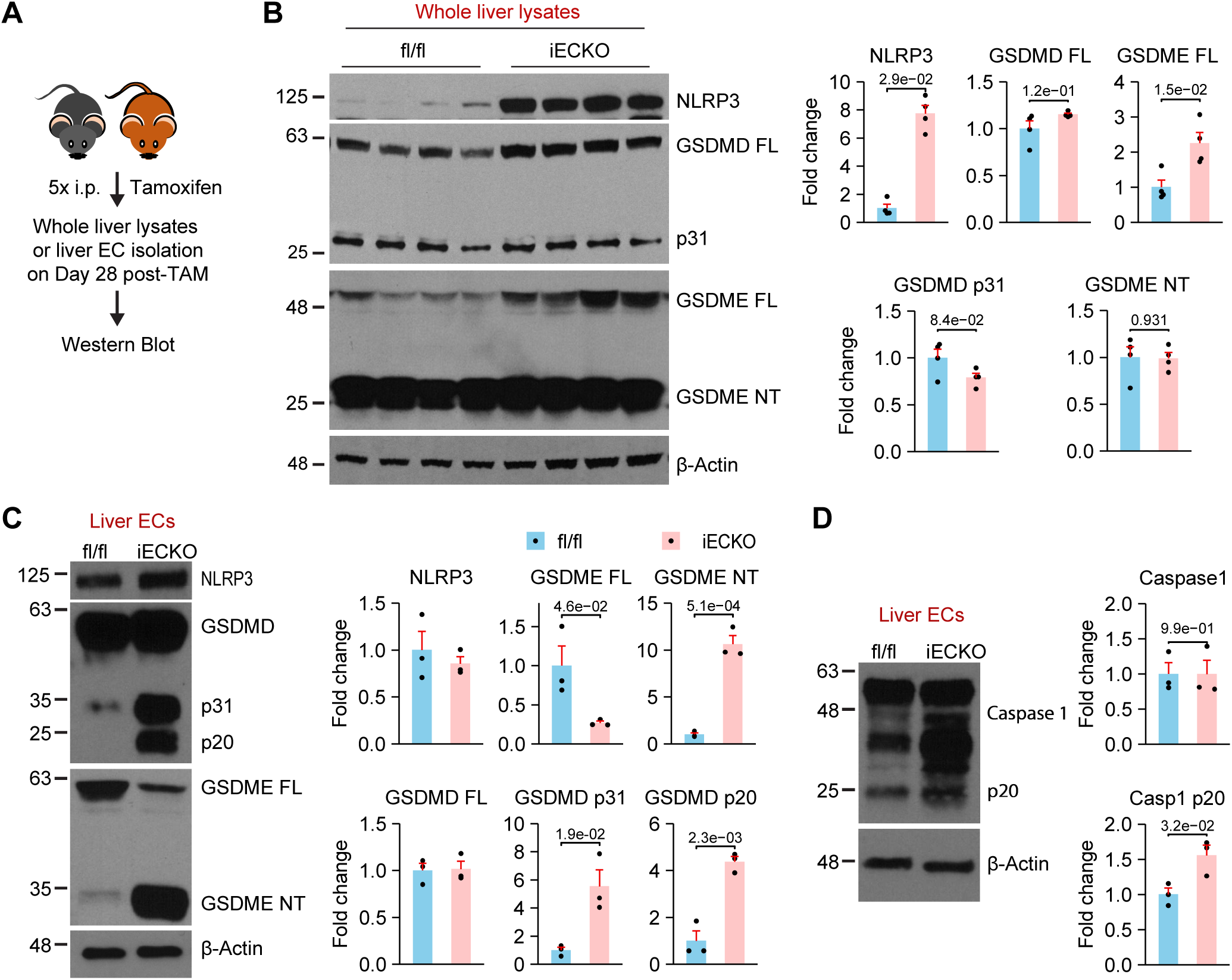
Endothelial neddylation deficiency activates gasdermin-mediated pyroptosis in the hepatic endothelium. **A**, Schematic depicting the workflow. The same liver lobes from *Nae1* fl/fl and iECKO mice were snap-frozen for storage, while the remaining tissue was used for EC isolation 28 days after tamoxifen administration. **B**, Protein expression and quantification in whole liver lysates assessed by Western Blot. n = 4 mice per group. **C and D**, Protein expression and quantification in isolated liver ECs assessed by Western Blot. Each point represents the data of ECs pooled from three livers. Statistical significance (*P* < 5.0e-2) was determined by the Mann-Whitney *U* test (NLRP3 in B) and the Student’s *t-*test (all other comparisons).

We therefore isolated CD31+ hepatic ECs for immunoblot analysis (Figure 5C-5D). In contrast to whole-liver lysates, *Nae1* deletion induced robust cleavage of both GSDMD and GSDME in hepatic ECs (Figure 5C), whereas total NLRP3 expression remained unchanged. Caspase-1 cleavage was also markedly increased (Figure 5D), consistent with inflammasome activation. These findings demonstrate that endothelial neddylation deficiency activates GSDMD/GSDME-dependent pyroptotic signaling selectively within the hepatic endothelium.

### Dual GSDMD/GSDME silencing reverses transcriptomic remodeling and attenuates liver injury induced by endothelial *Nae1* deletion

To determine whether GSDMD and GSDME mediate tissue injury caused by endothelial neddylation deficiency, we generated adeno-associated virus serotype 9 (AAV9) vectors expressing short hairpin RNAs (shRNAs) targeting both Gsdmd and Gsdme. Simultaneous silencing was chosen because both GSDMs were activated in *Nae1* iECKO ECs (Figure 5), and GSDME can compensate for GSDMD deficiency by mediating a salvage pyroptotic pathway^57^. *Nae1* fl/fl and *Nae1* iECKO mice received AAV9 expressing either control shRNA or dual GSDMD/GSDME shRNAs, followed by bulk RNA sequencing of liver tissue 26 days after *Nae1* deletion (Figure 6A).

**Figure 6.**
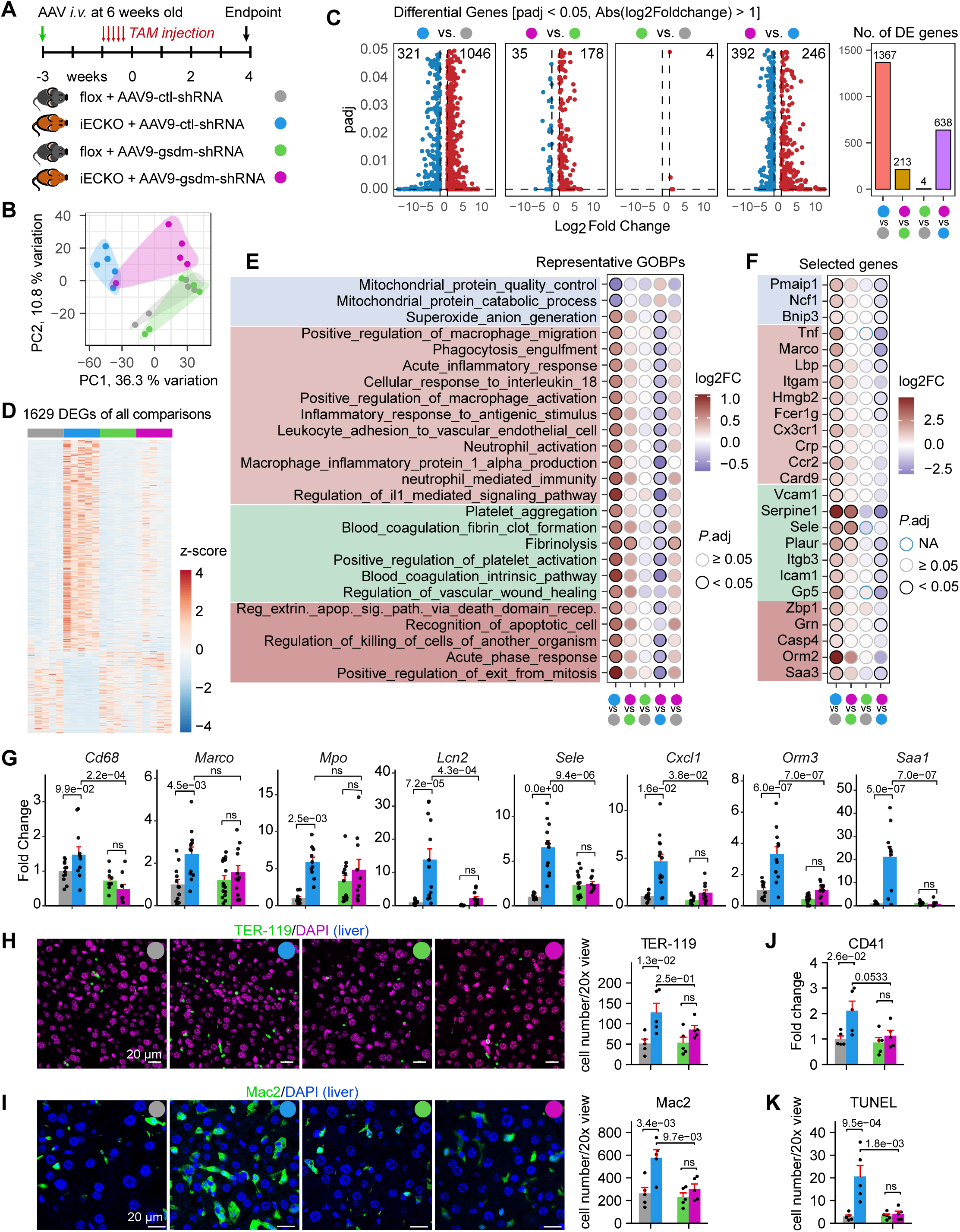
Dual GSDMD/GSDME silencing reverses transcriptomic remodeling and attenuates liver injury induced by endothelial *Nae1* deletion. **A**, Schematic representation of the experimental workflow for assessing GSMDs in mediating the observed phenotypic changes in iECKO mice. **B**, Principal component analysis reveals the global transcriptomic differences among the four groups of mice in A. Each dot represents an individual mouse (n=5 mice per group). **C**, Volcano plots of differentially expressed genes comparing the indicated groups and a bar plot showing the total number of DEGs in each comparison. **D**, A heat map displays z-scores of normalized expression levels of 1,629 DEGs (*P*adj < 0.05, |log2Foldchnage| > 1) for all comparisons among groups. **E**, Gene Set Variation Analysis identified up- or down-regulated Gene Ontology Biological Processes (GOBPs) in the five indicated comparisons. These GOBP terms are associated with mitochondrial dysfunction, inflammation & immunity, coagulation, cell death, and tissue damage. The color of circle edges reflects adjusted *P* value (*P*adj). Circles are filled with colors that represent the values of log2FC (log2foldchange) of GOBPs for each comparison. **F**, A dot plot showing the expression levels of selected genes involved in the GOBP pathways. The color of circle edges reflects adjusted *P* value. Circles are filled with colors that represent the values of log2FC (log2foldchange) of these genes for each comparison. **G**, The mRNA levels of the indicated genes assessed by qPCR in the livers of *Nae1* fl/fl and iECKO mice administered with AAV9-ctl-shRNA or AAV9-gsdmd/gsdme-shRNA. **H – K**, Immunostaining and/or quantification of TER-119 (H), Mac2 (I), J (CD41), and K (TUNEL) in the livers of the indicated mice. Statistical significance (*P* < 5.0e-2) was determined by the two-way ANOVA with Tukey’s HSD post-hoc test (G – K).

Principal component analysis demonstrated clear separation between *Nae1* fl/fl and Nae1 iECKO mice, with genotype representing the dominant source of transcriptomic variation (Figure 6B). Although dual GSDMD/GSDME silencing had minimal effects in *Nae1* fl/fl mice, it shifted the transcriptomic profile of *Nae1* iECKO mice toward that of control mice. Consistent with these findings, endothelial *Nae1* deletion induced 1,367 DEGs relative to control shRNA-treated *Nae1* fl/fl mice, whereas only 213 DEGs remained after dual GSDMD/GSDME silencing (Figure 6C). Likewise, dual GSDMD/GSDME silencing altered the expression of only four genes in *Nae1* fl/fl mice but reprogrammed 638 genes in *Nae1* iECKO mice relative to their control shRNA-treated counterparts. A heatmap of 1,629 DEGs detected across all groups further demonstrated broad normalization of the transcriptomic alterations induced by endothelial *Nae1* deletion (Figure 6D).

Gene Set Variation Analysis (GSVA) showed that *Nae1* deletion significantly enriched pathways associated with inflammation and innate immunity, including IL-1 signaling, leukocyte adhesion to vascular ECs, acute inflammatory responses, and neutrophil-mediated immunity (Figure 6E). Pathways related to platelet activation, coagulation, vascular wound healing, apoptotic cell recognition, acute-phase responses, and mitochondrial dysfunction were also significantly altered. These changes were largely abolished following dual GSDMD/GSDME silencing, with no significant pathway differences remaining between GSDMD/GSDME shRNA-treated *Nae1* iECKO and *Nae1* fl/fl mice. Expression patterns of representative genes closely paralleled the pathway-level changes identified by GSVA (Figure 6F).

Quantitative RT-PCR confirmed these transcriptomic findings (Figure 6G). Expression of inflammatory and tissue injury-associated genes, including *Cd68*, *Marco*, *Mpo*, *Lcn2*, *Sele*, *Cxcl1*, *Orm3*, and *Saa1*, was markedly increased in control shRNA-treated *Nae1* iECKO mice and substantially attenuated following dual GSDMD/GSDME silencing.

To determine whether transcriptomic rescue translated into improved tissue integrity, we examined vascular leakage, inflammation, cell death, and platelet accumulation in liver sections collected 26 days after *Nae1* deletion. As expected, control shRNA-treated *Nae1* iECKO mice exhibited increased TER-119-positive erythrocyte extravasation, Mac2-positive macrophage infiltration, TUNEL-positive cell death, and CD41-positive platelet accumulation compared with control *Nae1* fl/fl mice (Figures 6H–6K). Dual GSDMD/GSDME silencing markedly reduced each of these pathological features, demonstrating that GSDM activation is a major mediator of vascular injury and inflammation downstream of endothelial *Nae1* deletion.

Together, these findings identify GSDMD and GSDME as critical downstream effectors of endothelial neddylation deficiency. Simultaneous inhibition of both GSDMs substantially reverses transcriptomic remodeling, suppresses inflammatory responses, and mitigates vascular injury, hepatocyte death, immune cell infiltration, and platelet accumulation in *Nae1* iECKO mice.

### The neddylation pathway is downregulated in human vascular disease and experimental endotoxemia

To determine whether the neddylation pathway is altered in human vascular disease, we analyzed six publicly available Gene Expression Omnibus (GEO) datasets. Four bulk transcriptomic datasets included human carotid arteries with or without atherosclerotic lesions (GSE100927), early and advanced atherosclerotic plaques (GSE28829), carotid arteries containing atheroma plaques with paired macroscopically intact tissue (GSE43292), and blood samples from healthy individuals and patients with COVID-19 (GSE217948). Across these datasets, the expressions of *NAE1*, *CUL1*, and *CUL3* progressively declined with advancing atherosclerosis and were similarly reduced in blood cells from patients with COVID-19 (Figure 7A).

**Figure 7.**
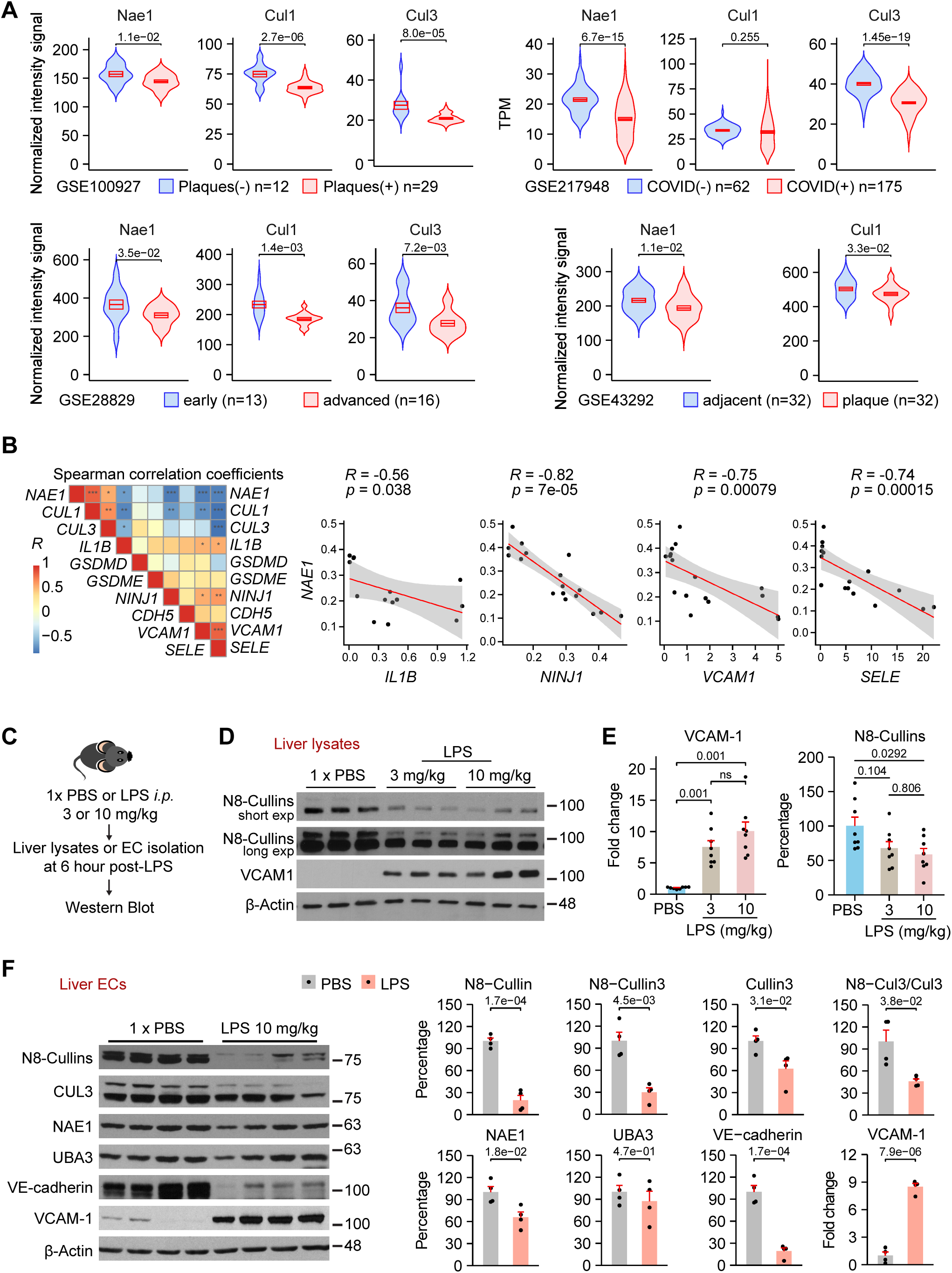
The neddylation pathway is downregulated in human vascular disease and experimental endotoxemia. **A**, The mRNA expression levels of *NAE1*, *CUL1*, *CUL3* in public datasets GSE100927, GSE217948, GSE28829, and GSE43292 were shown. The top and bottom edges of boxes represent the standard errors of the means, and the middle lines of boxes represent the means. **B**, The heatmap displays Spearman’s correlation coefficients for the indicated gene expression levels in the EC populations, utilizing 19 samples from two human arterial scRNA-seq datasets (GSE131778 and GSE198750). **C**, Experimental workflow for D – F. **D** and **E**, Western Blot and quantification of the indicated proteins in whole liver lysates. Total 24 mice were used (n = 8 mice per group). **F**, Western Blot of the indicated proteins in the isolated liver ECs. Each sample represents ECs from three livers. Total 24 mice were used. Statistical significance (*P* < 5.0e-2) was determined by Student’s *t-*test (*NAE1* and *CUL1* in GSE100927, *NAE1* and *CUL3* in GSE28829, *NAE1* and *CUL1* in GSE43292, and comparisons except N8-CUL3/CUL3 in F), Welch’s *t*-test (*CUL1* in GSE28829 and N8-CUL3/CUL3 in F), Mann-Whitney *U* test (*CUL3* in GSE100927 and all genes in GSE217948), Welch’s Anova followed by Games-Howell post-hoc test (VCAM-1 in E), and one-way Anova followed by Tukey HSD post-hoc test (Neddylation in E).

To further examine endothelial-specific changes, we analyzed two human arterial single-cell RNA-sequencing datasets (GSE131778 and GSE198750), comprising 7,631 ECs among nine major cell populations (Figure S5). Within ECs, *NAE1* expression was inversely correlated with *NINJ1*, *VCAM1*, *SELE*, and *IL1B* (Figure 7B), linking reduced neddylation with endothelial activation and inflammatory dysfunction in human vascular disease.

Because endothelial neddylation deficiency recapitulated several features of endotoxin-induced endothelial dysfunction, including inflammatory activation, increased adhesion molecule expression, and a procoagulant phenotype, we next examined the neddylation pathway in lipopolysaccharide (LPS)-treated mice (Figure 7C). Immunoblot analysis of whole-liver lysates demonstrated increased VCAM-1 expression accompanied by reduced levels of neddylated cullins following LPS administration (Figures 7D, 7E, and S6). Similar changes were observed in isolated CD31+ hepatic ECs, which exhibited increased VCAM-1 expression, reduced VE-cadherin, and marked loss of neddylated cullins, including neddylated Cullin-3 (Figure 7F). In contrast, expression of the NEDD8-activating enzyme subunits NAE1 and UBA3 was modestly reduced, consistent with the diminished neddylation induced by LPS.

Together, these findings demonstrate that endothelial neddylation is broadly suppressed during both chronic vascular disease and acute inflammatory injury.

### Restoration of endothelial neddylation partially reverses LPS-induced endothelial transcriptomic remodeling

Because neddylation was suppressed in hepatic ECs following LPS treatment, we next tested whether restoring endothelial neddylation could mitigate the inflammatory transcriptional response. We generated adeno-associated virus serotype 9 (AAV9) vectors expressing the neddylation machinery, including the E1 enzyme subunits NAE1 and UBA3 together with the E2 enzymes UBE2M and UBE2F, under control of the endothelial-specific ICAM2 promoter. C57BL/6 mice received intravenous administration of AAV9 expressing either GFP or E1/E2 enzymes, followed by LPS challenge and bulk RNA sequencing of isolated hepatic ECs (Figure 8A).

**Figure 8.**
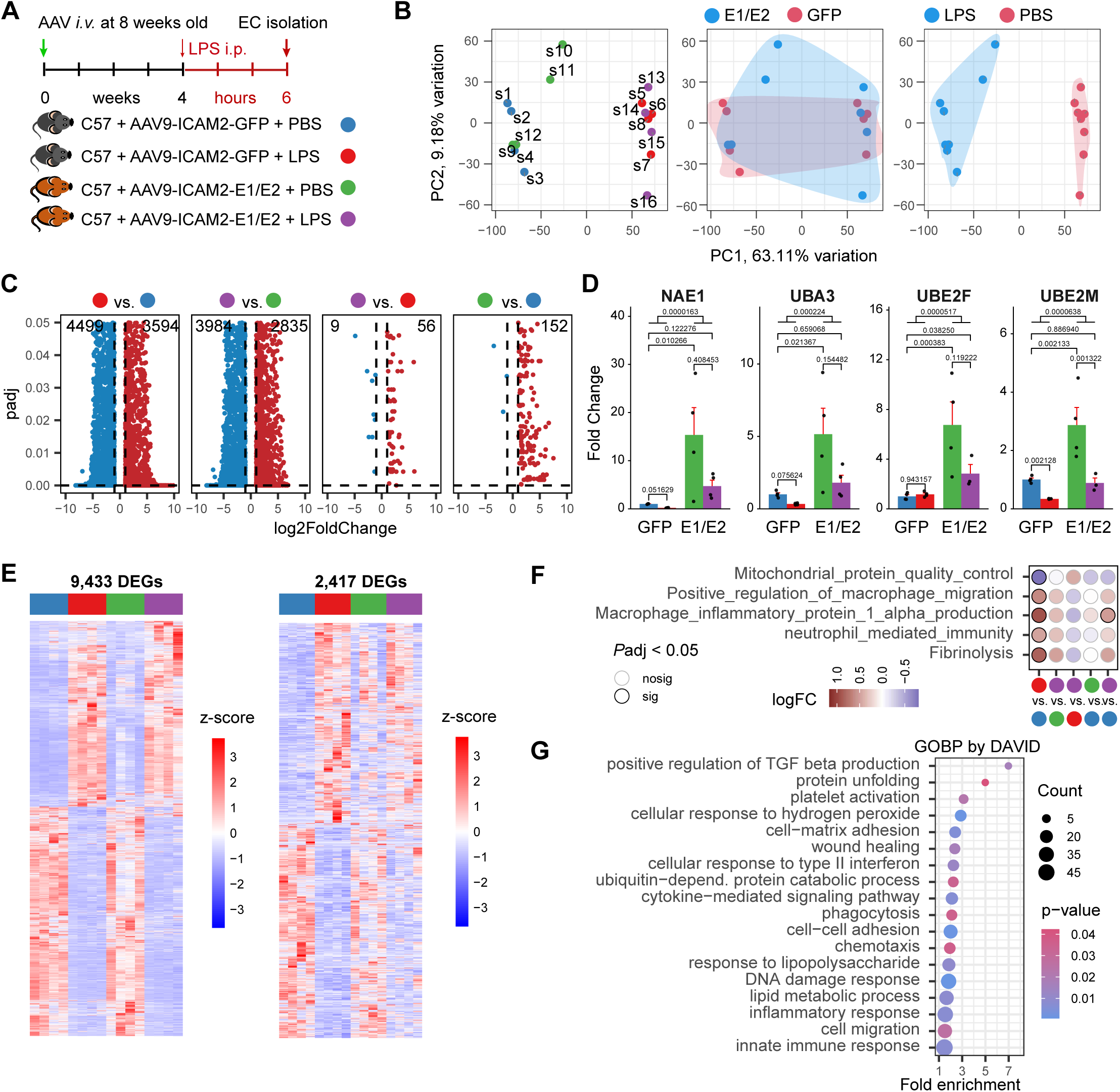
Restoration of endothelial neddylation partially reverses LPS-induced endothelial transcriptomic remodeling. **A**, Experimental workflow. Liver ECs were isolated from 48 mice (12 mice per group). The total RNAs after ribosomal RNAs depletion were used for bulk RNA-seq analysis. **B**, Principal component analysis reveals the global transcriptomic differences among the four groups of mice in A. Each dot represents ECs from three livers. **C**, Volcano plots display DEGs comparing the indicated groups and a bar plot shows the total number of DEGs in each comparison. **D**, The mRNA expression levels of neddylation E1 (*Nae1* and *Uba3*) and E2 (*Ube2f* and *Ube2m*) enzymes were examined in the isolated ECs. **E**, Heatmaps display z-scores of normalized expression levels of 9,433 DEGs (*P*adj < 0.05, |log2Foldchnage| > 1) for all comparisons among groups and 2417 DEGs (*P*adj < 0.05, |log2Foldchnage| > 0.585) that are differentially expressed between LPS and PBS treatments in GFP-expressing mice, but not in neddylation E1/E2-expressing mice. **F**, Gene Set Variation Analysis identified up- or down-regulated Gene Ontology Biological Processes (GOBPs) in the LPS vs. PBS comparison in GFP-expressing mice but not in neddylation E1/E2-expressing mice. The color of circle edges reflects adjusted *P* value. Circles are filled with colors that represent the values of log2FC (log2foldchange) of GOBPs for each comparison. **G**, A dot plot illustrates the enriched GOBP terms associated with the 2,417 DEGs identified by DAVID. Statistical significance (*P* < 5.0e-2) was determined by the two-way ANOVA with Tukey’s HSD post-hoc test after log-transformation.

Principal component analysis demonstrated that LPS was the dominant determinant of transcriptomic variation, with the first two principal components accounting for 63.1% and 9.2% of the total variance, respectively (Figure 8B). Consistent with this observation, LPS induced extensive transcriptional remodeling, with 8,093 DEGs identified relative to PBS-treated GFP controls (Figure 8C). Restoration of endothelial neddylation reduced this number to 6,810 DEGs, whereas E1/E2 expression alone produced minimal transcriptomic changes under either basal or inflammatory conditions, altering only 152 genes in PBS-treated mice and 65 genes in LPS-treated mice. Quantitative RT-PCR confirmed robust endothelial transgene expression, with E1/E2 transcripts increased by as much as 15-fold (Figure 8D).

Among the genes altered by LPS, 2,417 were no longer differentially expressed after restoration of endothelial neddylation (Figure 8E), indicating partial normalization of the inflammatory transcriptional program. Gene Set Variation Analysis further demonstrated that LPS enriched pathways associated with macrophage migration, macrophage inflammatory protein-1α production, neutrophil-mediated immunity, and fibrinolysis while suppressing mitochondrial protein quality control (Figure 8F). These pathway alterations were largely reversed by E1/E2 expression. Functional enrichment analysis of the rescued genes further identified inflammatory responses, immune regulation, extracellular matrix remodeling, and protein homeostasis as the principal biological processes restored by enhanced neddylation (Figures 8G and S7).

Cullins are the predominant neddylated proteins in ECs, and their neddylation was markedly reduced following LPS treatment (Figure 7D and 7F). To determine whether diminished cullin neddylation contributes to LPS-induced EC injury, we treated mice with CSN5i-3 ^58^, a selective inhibitor of the COP9 signalosome (CSN), the principal deneddylase for cullins^22,23^, prior to LPS administration to preserve cullin neddylation (Figure S8A). Transcriptomic analysis of hepatic ECs revealed that CSN5i-3 markedly attenuated LPS-induced transcriptional remodeling, reducing the number of differentially expressed genes (DEGs) relative to control from 9,279 in the LPS group to 6,742 in the CSN5i-3+LPS group (Figures S8B and S8C). Among the genes dysregulated by LPS, 4,383 were restored toward control levels by CSN5i-3 treatment (Figure S8D). Gene Set Variation Analysis further showed that multiple pathways induced by LPS, including cellular responses to interleukin-18, macrophage activation, interleukin-1 signaling, and the intrinsic coagulation pathway, were no longer significantly enriched following CSN5i-3 treatment (Figure S8E). Notably, enrichment of the pyroptotic cell death pathway was also largely abolished by CSN5i-3 (Figure S8F).

Together, these findings demonstrate that suppression of endothelial neddylation is a feature of both human vascular disease and experimental inflammation and that restoring neddylation partially normalizes inflammatory and extracellular matrix remodeling programs in hepatic ECs.

## Discussion

The present study identifies NAE1-mediated protein neddylation as a fundamental regulator of endothelial homeostasis and vascular integrity. Using an inducible endothelial-specific *Nae1* deletion model, we demonstrate that disruption of endothelial neddylation results in catastrophic vascular failure characterized by loss of endothelial identity, vascular leakage, platelet accumulation, inflammatory cell recruitment, and multi-organ injury (Figures 1 and 3). Integrated transcriptomic, proteomic, and biochemical analyses further identify GSDM-dependent pyroptosis as a major mechanism linking loss of endothelial neddylation to endothelial dysfunction (Figures 2, 4, 5). Mechanistically, endothelial neddylation deficiency activated both GSDMD- and GSDME-dependent pyroptotic pathways, whereas simultaneous inhibition of GSDMD and GSDME substantially attenuated the transcriptional and pathological consequences of *Nae1* deficiency (Figure 6). Finally, we demonstrate that endothelial neddylation is suppressed in both experimental inflammation and human vascular disease and that restoration of the neddylation pathway partially reverses inflammatory endothelial reprogramming (Figures 7 and 8). Collectively, these findings establish endothelial protein neddylation as a previously unrecognized regulator of vascular homeostasis and identify this pathway as a potential therapeutic target in inflammatory vascular disease.

A central finding of this study is that constitutive protein neddylation represents a previously unrecognized post-translational mechanism for maintaining endothelial identity under physiological conditions. Although endothelial activation is well-recognized hallmark of cardiovascular and inflammatory diseases^1,2,9^, the post-translational mechanisms that preserve endothelial identity and vascular homeostasis, particularly those mediated by protein neddylation, have remained largely unexplored. Here, we demonstrate that constitutive neddylation is indispensable for endothelial homeostasis. Loss of *Nae1* induced broad transcriptomic and proteomic reprogramming, shifting ECs from a quiescent vascular program toward inflammatory, procoagulant, and mesenchymal-like phenotypes that culminated in vascular leakage, hemorrhage, thrombosis, and multi-organ failure.

Among the expanding repertoire of NEDD8 substrates, cullins are the predominant NEDD8 targets in ECs, and robust cullin neddylation are readily detected under physiological conditions. Neddylation promotes the assembly and activation of CRLs, which maintain protein homeostasis through ubiquitin-dependent degradation of numerous signaling proteins^59^. Although CRLs have long been regarded as the principal mediators of neddylation-dependent biology, their role in maintaining endothelial identity and vascular homeostasis has remained poorly defined. Given the fundamental role of neddylation in preserving endothelial quiescence and preventing spontaneous transition toward pathological activation, future studies should systematically define the endothelial NEDD8 proteome and determine how neddylated cullins and non-cullin substrates cooperatively maintain endothelial identity and vascular homeostasis.

A major mechanistic advance of this study is the identification of GSDMD- and GSDME-mediated pyroptosis as a key downstream effector of endothelial dysfunction following loss of neddylation. Although pyroptosis has emerged as an important contributor to atherosclerosis^62,63^, myocardial infarction^64^, ischemia-reperfusion injury^65^, sepsis^66^, and other cardiovascular disorders, previous studies have largely focused on immune cells as the predominant source of pyroptotic signaling. Here, we demonstrate that neddylation functions as an endogenous safeguard of endothelial survival by restraining GSDMD- and GSDME-dependent pyroptosis under physiological conditions. Single-cell transcriptomic analyses demonstrated progressive enrichment of pyroptotic programs before widespread endothelial loss, while biochemical analyses confirmed activation of both GSDMD and GSDME in hepatic ECs. Importantly, simultaneous silencing of GSDMD and GSDME substantially attenuated inflammatory gene expression, vascular leakage, platelet accumulation, hepatocyte injury, and immune-cell infiltration, indicating that GSDM activation is not merely a consequence of endothelial injury but a major driver of disease progression. Additionally, our findings also expand the biological functions of neddylation beyond its well-established role in apoptosis^60^ by identifying suppression of pyroptosis as a critical mechanism through which neddylation maintains vascular integrity. Although additional forms of regulated cell death may also contribute to disease progression, our findings support a model in which constitutive endothelial neddylation maintains vascular integrity by suppressing GSDM activation, whereas loss of neddylation lowers the threshold for endothelial pyroptosis, amplifying inflammation and propagating tissue injury through disruption of the vascular barrier.

Our findings further establish the clinical relevance of endothelial neddylation by demonstrating that this pathway is dynamically regulated during inflammatory vascular injury. Analysis of multiple human transcriptomic datasets revealed reduced expression of key components of the neddylation machinery in atherosclerotic arteries and inflammatory diseases, including COVID-19, while experimental endotoxemia similarly suppressed endothelial neddylation in mice. These observations suggest that impaired endothelial neddylation is a conserved feature of inflammatory vascular disorders rather than a phenomenon unique to genetic *Nae1* deficiency. Pharmacological inhibition of neddylation with MLN4924 has demonstrated promising antitumor activity^60^ but is associated with dose-limiting toxicities, including hepatotoxicity, neutropenia, and anemia ^68,69^. Given the severe vascular injury caused by endothelial neddylation deficiency in our study, endothelial dysfunction may contribute to some of these adverse effects. These findings therefore underscore the need to carefully evaluate endothelial and vascular toxicity during systemic neddylation inhibition and highlight the importance of dose, duration, and tissue specificity when targeting this pathway therapeutically.

Lastly, our study provides proof-of-concept that restoring endothelial neddylation mitigates inflammatory vascular injury. In an endotoxemia model, enhancement of endothelial neddylation using two complementary approaches—genetic augmentation of the neddylation pathway and pharmacological inhibition of cullin deneddylation—substantially reversed LPS-induced endothelial transcriptional reprogramming, particularly pathways associated with inflammation, extracellular matrix remodeling, leukocyte recruitment, pyroptosis, and mitochondrial dysfunction. These findings suggest that inflammatory stimuli suppress endothelial neddylation, thereby promoting endothelial pyroptosis, disrupting vascular homeostasis, and exacerbating vascular injury. More importantly, they identify restoration of endothelial neddylation as a conceptually distinct therapeutic strategy that targets a fundamental homeostatic mechanism rather than individual downstream inflammatory pathways. Future studies should determine whether endothelial-specific enhancement of neddylation improves vascular function in chronic cardiovascular diseases, elucidate the mechanisms by which inflammatory stimuli impair endothelial neddylation, and develop endothelial-targeted therapeutic approaches that maximize vascular protection while minimizing systemic toxicity.

Several limitations should be considered. First, our mechanistic studies were performed predominantly in murine models and cultured ECs. Although analyses of multiple human transcriptomic datasets support the clinical relevance of our findings, direct assessment of endothelial neddylation activity in human vascular tissues will be important. Second, although our data identify GSDMD and GSDME as major downstream mediators of endothelial dysfunction, the molecular mechanisms linking impaired neddylation to GSDM activation remain undefined. Our transcriptomic and proteomic analyses consistently demonstrated activation of NRF2-dependent signaling following *Nae1* deletion, consistent with impaired Cullin-3-dependent ubiquitination and confirming effective disruption of CRL activity. Whether GSDMs or their upstream regulators are directly controlled by neddylation or are activated indirectly through altered ubiquitin-dependent signaling requires further investigation. Third, although GSDM inhibition substantially ameliorated the phenotype, additional mechanisms—including endothelial-to-mesenchymal transition, oxidative stress, and extracellular matrix remodeling—are also likely to contribute to endothelial dysfunction following loss of neddylation. Finally, our study focused primarily on the liver and lung vasculature, and whether similar mechanisms operate across distinct endothelial subtypes and vascular beds remains to be determined.

In conclusion, this study identifies protein neddylation as a fundamental post-translational mechanism that preserves endothelial identity, vascular integrity, and tissue homeostasis. Constitutive endothelial neddylation maintains vascular quiescence through continuous CRL-dependent protein turnover and suppression of GSDM-mediated pyroptosis. Loss of this pathway triggers endothelial phenotypic reprogramming, vascular barrier disruption, thrombosis, and multiorgan injury, whereas restoration of neddylation mitigates inflammatory endothelial remodeling. Together, these findings establish endothelial protein neddylation as a central regulator of vascular homeostasis and support therapeutic modulation of this pathway as a promising strategy for inflammatory vascular disease.

## Nonstandard Abbreviations and Acronyms

AAV: adeno-associated virus
ECs: endothelial cells
CRLs: Cullin-RING E3 ubiquitin ligases
CSN: the COP9 signalosome
DEGs: differentially expressed genes
FITC: fluorescein isothiocyanate
fl/fl: flox / flox
GOBP: Gene ontology biological process
GSDMD: gasdermin D
GSDME: gasdermin E
GSEA: gene set enrichment analysis
GSVA: gene set variation analysis
HUVEC: human umbilical vein endothelial cell
iECKO: inducible endothelial-cell-specific knockout
OCT: Optimal Cutting Temperature
LPS: lipopolysaccharide
MLN: MLN4924 (Pevonedistat)
NAE1: NEDD8 activating enzyme E1 subunit 1
NEDD8: neural precursor cell expressed developmentally downregulated 8
NRF2: nuclear factor erythroid 2-related factor 2
RNA-seq: RNA-sequencing
scRNA-seq: single-cell RNA-sequencing
shRNAs: short hairpin RNAs
snRNA-seq: single-nucleus RNA-sequencing
SUMO: small ubiquitin-like modifier
TER-119: transplantation erythroid record-119
TMT: Tandem Mass Tag
TUNEL: terminal deoxynucleotidyl transferase dUTP Nick-End Labeling
UBA3: ubiquitin like modifier activating enzyme 3
UMAP: Uniform Manifold Approximation and Projection
VCAM-1: vascular cell adhesion molecule-1

## Acknowledgments

The authors thank Dirk Anderson, Director, Flow Cytometry and Single-Cell Genomics Core Facilities at UNL, for generating the libraries of single-nucleus RNA-seq; Sandra Thibivilliers and Marc Libault for generating the libraries of single-cell RNA-seq or providing technical support. The authors thank the Proteomics and Metabolomics Facility at UNL for performing TMT10plex Isobaric Mass Tagging and mass spectrometry (Michael Naldrett and Sophie Alvarez). The authors also thank the Nebraska Veterinary Diagnostic Center at UNL for performing tissue sectioning and H&E staining. The authors acknowledge the Light & Electron Microscopy Core at UNL, the Nebraska Center for Integrated Biomolecular Communication, and its funding sources (National Institutes of Health National Institutes of General Medical Sciences P20GM113126). The authors also acknowledge the University of Kansas Medical Center Genomics Core for single-nucleus RNA-sequencing and data de-multiplexing and transfer and the funding source for the Core – Kansas Intellectual and Developmental Disabilities Research Center (NIH U54 HD090216), the Molecular Regulation of Cell Development and Differentiation – COBRE (P30 GM122731-03), the NIH S10 High-End Instrumentation Grant (NIH S10OD021743), and the Frontiers CTSA grant (UL1TR002366) at the University of Kansas Medical Center. Lastly, we thank Dr. Peng Xiao at the University of Nebraska Medical Center who helped with the Ingenuity Pathway Analysis. Some illustrations shown in this article were created using Biorender (BioRender.com).

## Author Contributions

R.N. Zoni, J. Li, H. Su, and X. Sun conceived the study, and R.N. Zoni, F. Tanni, and X. Sun wrote the article. R.N. Zoni, F. Tanni, C. Lee, X. Cheng, J.Y. Zhu, and X. Sun planned the experiments. R.N. Zoni, F. Tanni, C. Lee, X. Cheng, V.B. Baki, M.S. Islam, and J.Y. Zhu performed the experiments. X. Sun analyzed RNA sequencing and bioinformatic data. H. Su and X. Sun provided intellectual input. H. Su and X. Sun supervised the project. All authors read and approved the final paper.

## Supplementary Materials

### Supplementary Figure legends

**Figure S1.**
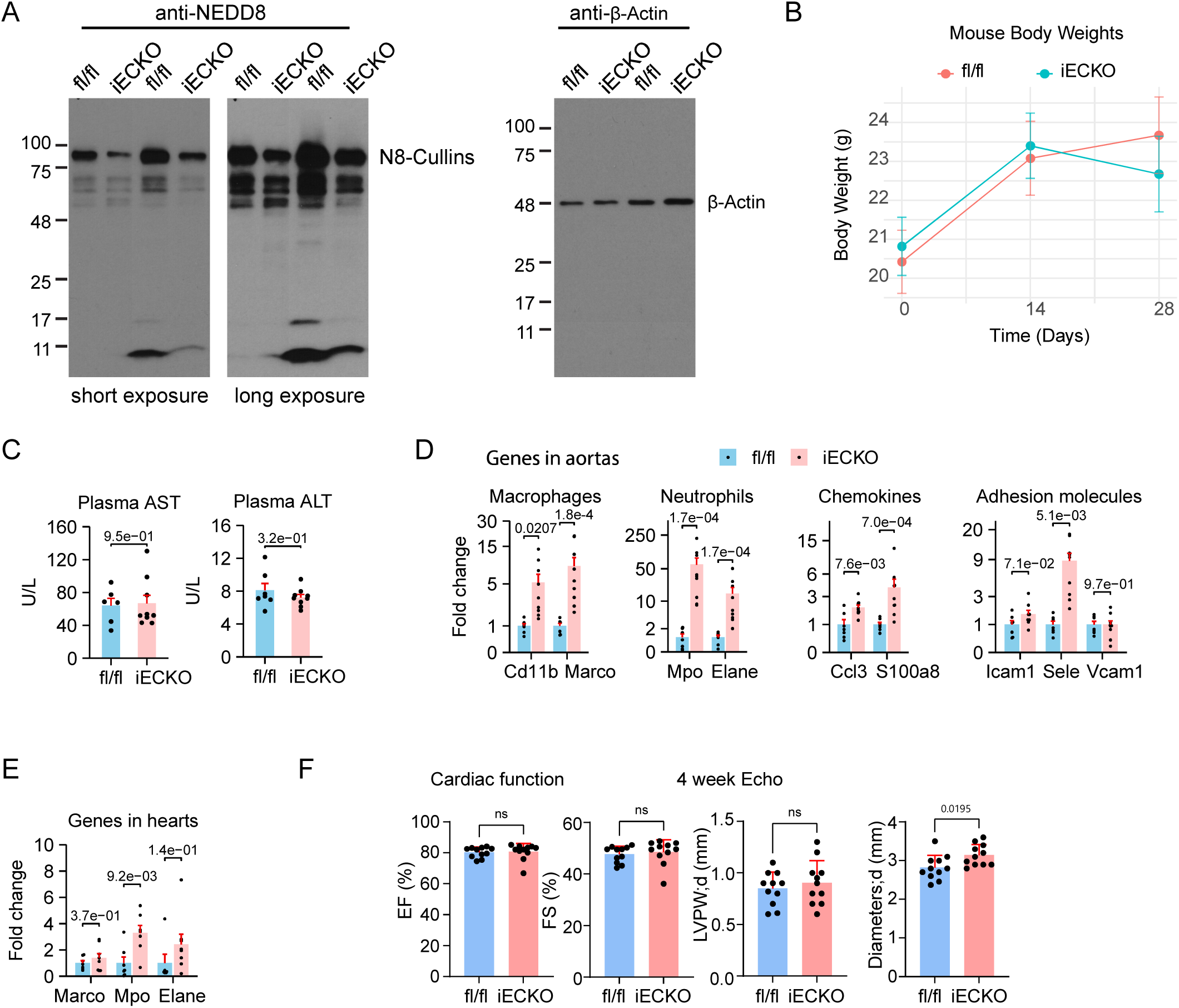
Molecular and physiological phenotypes associated with *Nae1* deletion in the vascular endothelium. **A**, Western blot analysis of neddylation in isolated liver ECs from *Nae1* fl/fl and iECKO mice 28 days after tamoxifen administration. **B**, Body weights of *Nae1* fl/fl and iECKO mice on the same day of tamoxifen administration and 14 or 28 days after tamoxifen administration. **C**, Plasma levels of AST and ALT 28 days after tamoxifen administration. **D**, The mRNA levels of the indicated genes assessed by qPCR in the aortas of *Nae1* fl/fl and iECKO mice 28 days after tamoxifen administration. **E**, The mRNA levels of the indicated genes assessed by qPCR in the hearts of *Nae1* fl/fl and iECKO mice 28 days after tamoxifen administration. **F**, Echocardiography of *Nae1* fl/fl and iECKO mice 28 days after tamoxifen administration. Statistical significance (*P* < 5.0e-2) was determined by Student’s *t-*test (C, *Ccl3* and *Vcam1* in D, *Marco* and *Mpo* in E, and F), Welch’s *t*-test (*Cd11b* and *Sele* in D), and Mann-Whitney *U* test (*Marco*, *Mpo*, *Elane*, and *Icam1* in D; *Elane* in E).

**Figure S2.**
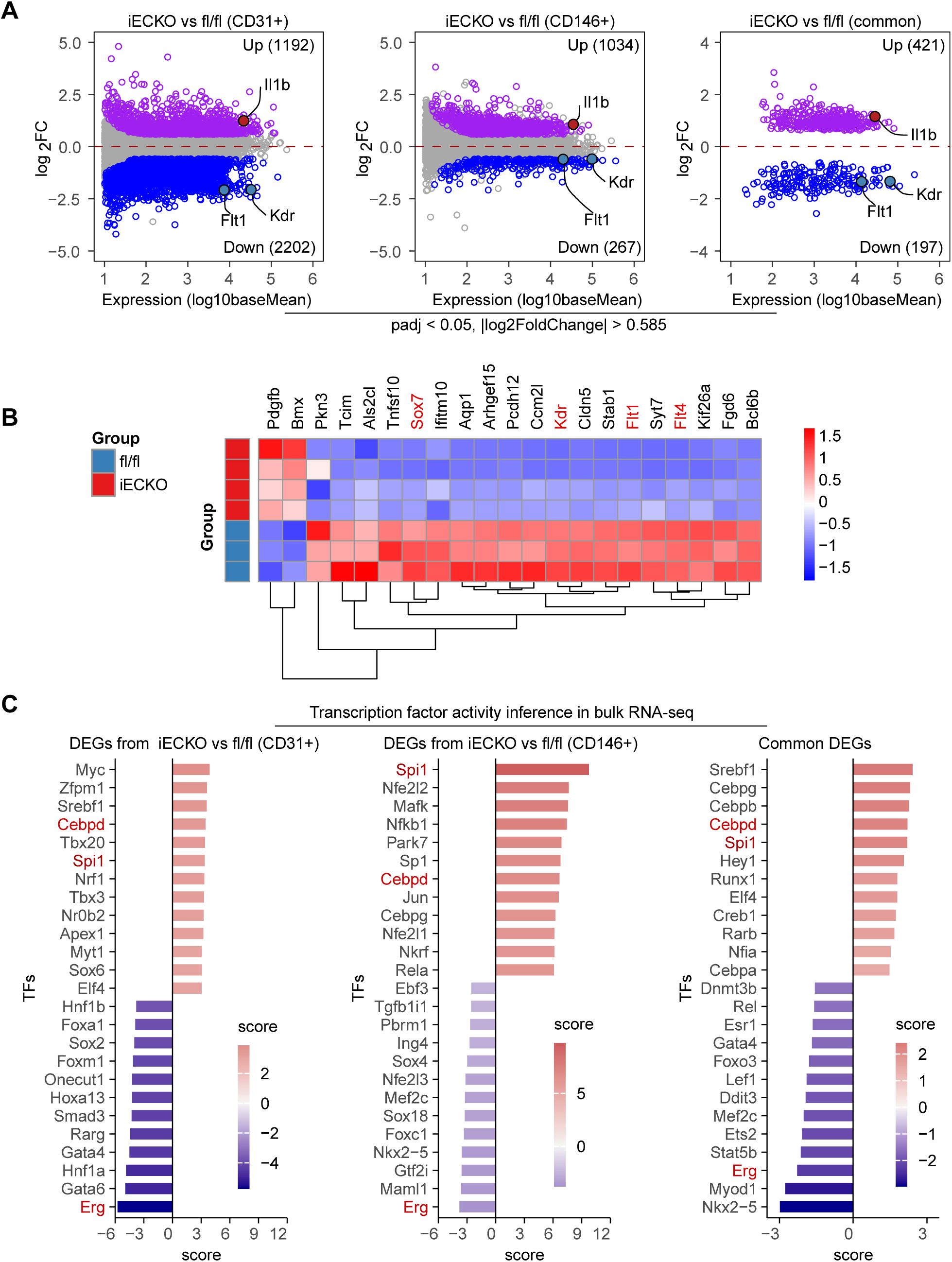
Bulk RNA-seq analysis of CD31+ or CD146+ cells isolated from fl/fl and iECKO mice 14 days after tamoxifen administration. **A**, MA plots display the DEGs (padj < 0.05, |log2FoldChange| > 0.585). **B**, A heatmap displays the expression levels of 21 genes overlapping between the Vascular EC Core Gene Set (GSE210119) and the 618 common DEGs. **C**, The transcription factor activities were inferred using the package decoupleR in R. Normalized and log transformed counts were used for decoupleR analysis.

**Figure S3.**
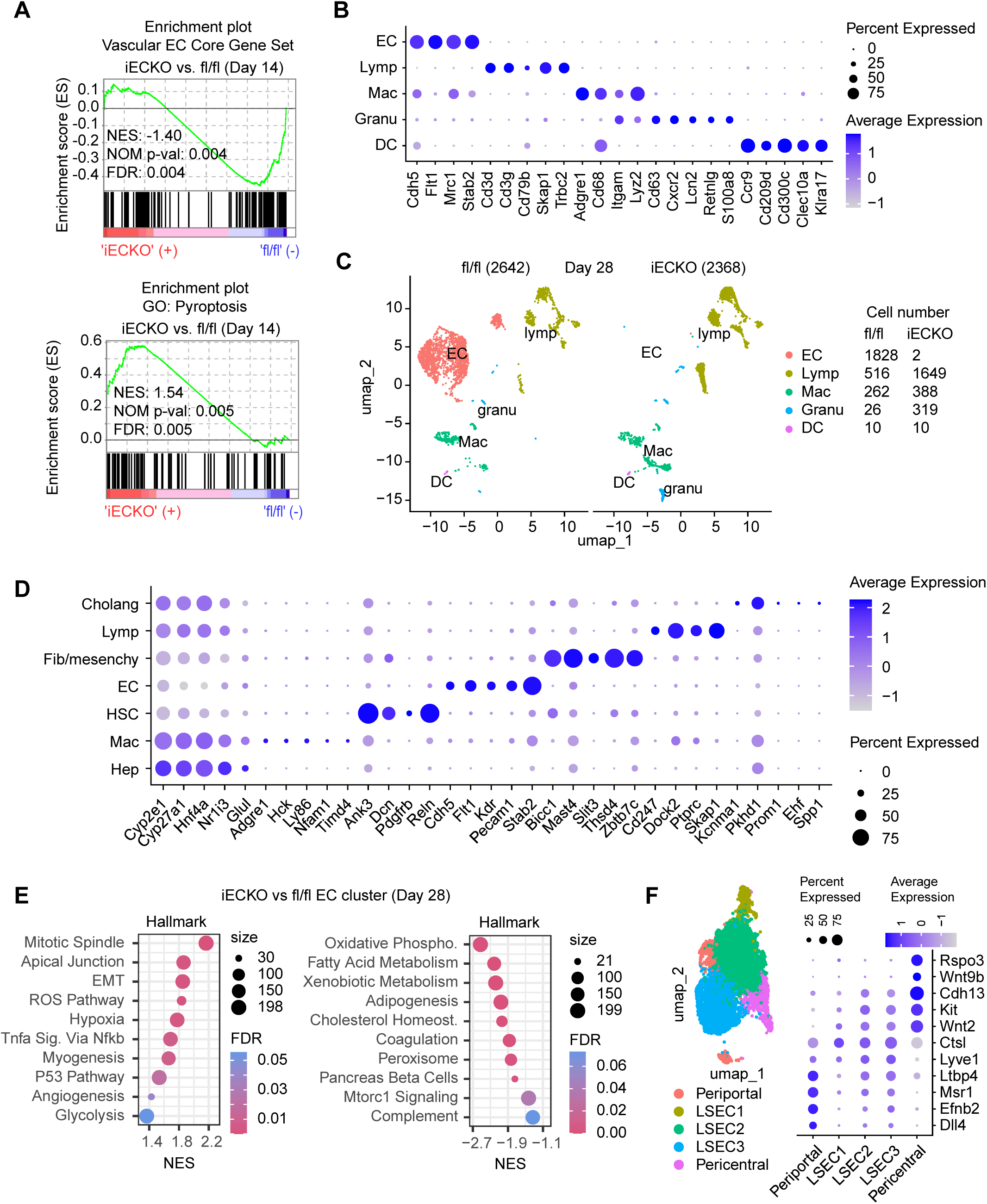
scRNA-seq analysis of CD31+ cells of livers from fl/fl and iECKO mice and snRNA-seq analysis of isolated nuclei of livers from fl/fl and iECKO mice. **A**, Enrichment plots showing that the Vascular EC Core Gene Set is downregulated and GO pyroptosis is upregulated in the EC cluster of iECKO sample at day 14 post tamoxifen injection. **B**, A dot plot showing the marker genes of annotated cell clusters identified by scRNA-seq. **C**, Uniform manifold approximation and projection (UMAP) of scRNA-seq for the *Nae1* day28 fl/fl (n=2,642 cells) and iECKO (n=2,368 cells) samples after quantity control and data filtering using Harmony integration. Cell numbers of annotated cell clusters in the fl/fl and iECKO samples were shown. **D**, A dot plot showing the marker genes of annotated cell clusters identified by snRNA-seq. **E**, GSEA revealed the enriched Hallmark pathways associated with the DEGs identified by snRNA-seq in the EC populations between fl/fl and iECKO mice 28 days after tamoxifen administration. **F**, EC subclusters 0, 1, 2, 3, and 4 were annotated as liver sinusoidal endothelial cells (LSEC) subcluster 2 (LSEC2), LSEC3, pericentral ECs, periportal ECs, and LSEC1. Lineage 1 progresses through distinct EC subpopulations including LSEC1, LSEC2, LSEC3, and periportal ECs, while lineage 2 progresses through LSEC1, LSEC2, and pericentral ECs.

**Figure S4.**
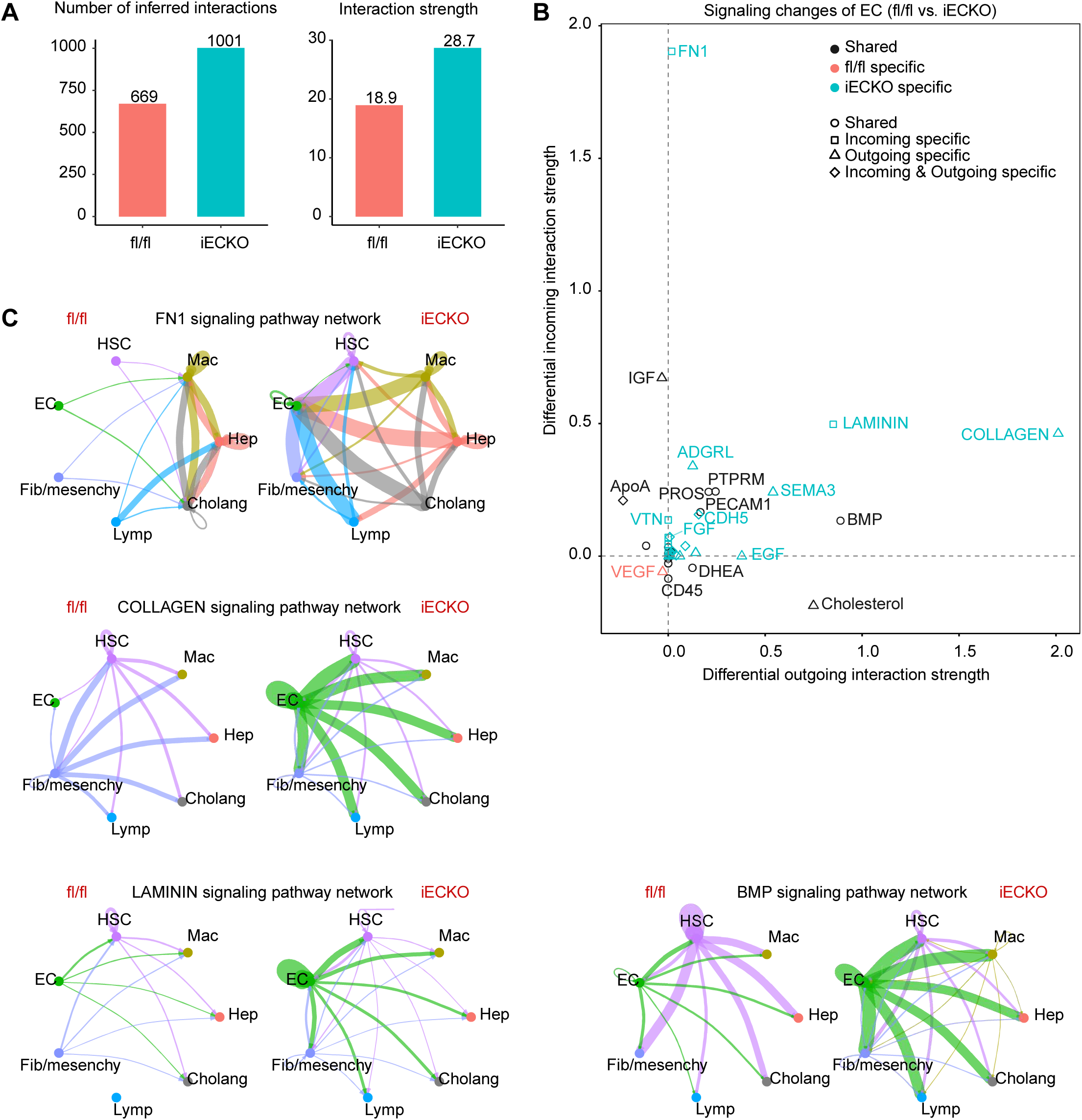
Cell-cell communications revealed by CellChat analysis of snRNA-seq data. **A**, Bar plots display the interaction numbers and strength in the fl/fl and iECKO livers. **B**, A differential incoming and outgoing interaction strength plot from CellChat analysis. A positive value at the x-axis means the iECKO ECs send more signals; a negative value means they send fewer signals. A positive value at the y-axis means iECKO ECs receive more signals; a negative value means they receive fewer signals. **C**, Circle plots display the indicated signaling pathway networks. A line looping back into the same cell group shows autocrine signaling (e.g., the green line in ECs). Line colors match the sending (source) cell group. For example, the green line indicates that the signaling is from ECs. The line width represents the strength of interactions between two cell groups.

**Figure S5.**
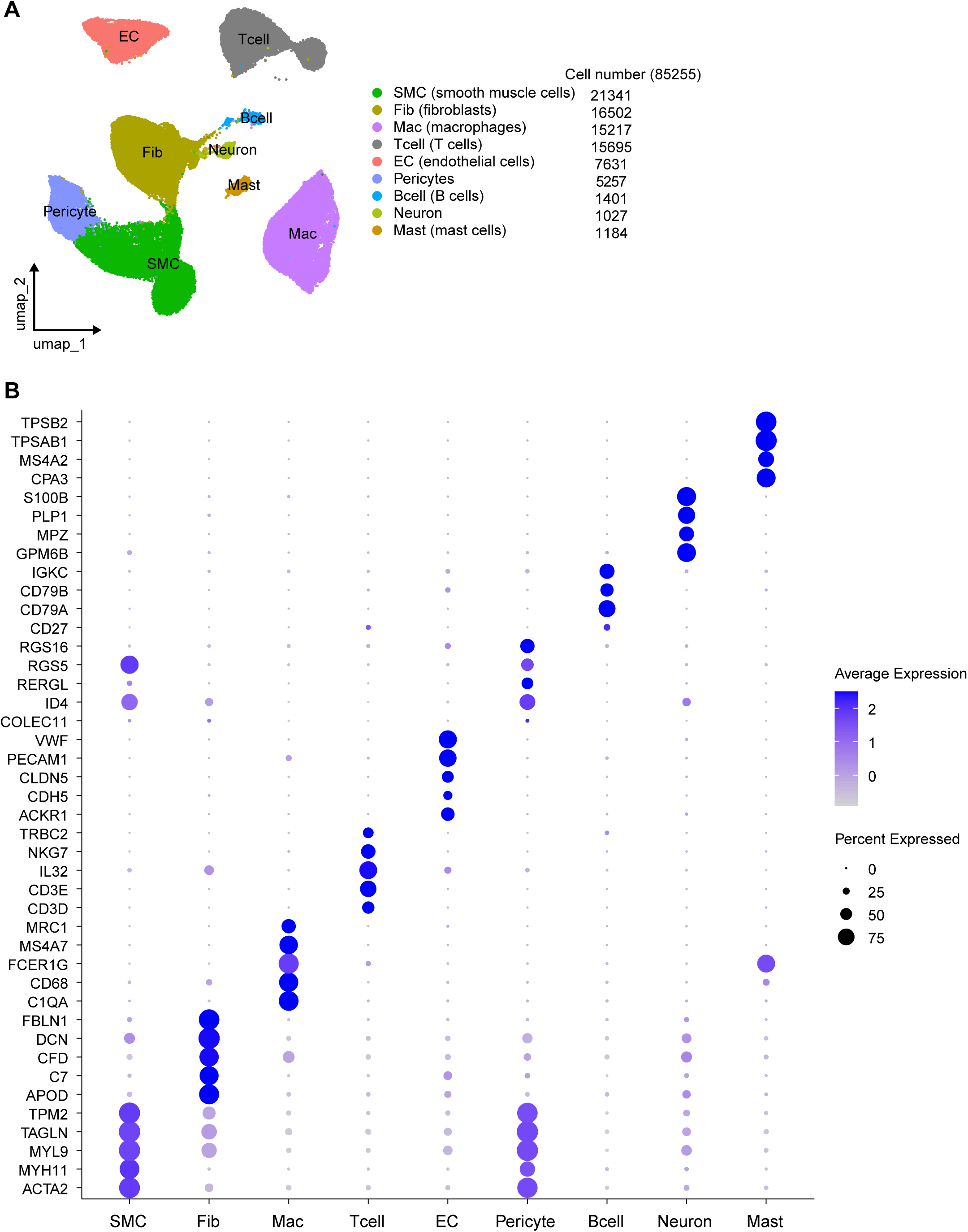
Analysis of human arterial scRNA-seq datasets (GSE131778 and GSE198750). **A**, A Uniform manifold approximation and projection (UMAP) of scRNA-seq datasets (GSE131778 and GSE198750). The number of cells was shown for the nine cell populations. **B**, A dot plot showing the marker genes of annotated cell clusters identified by scRNA-seq.

**Figure S6.**
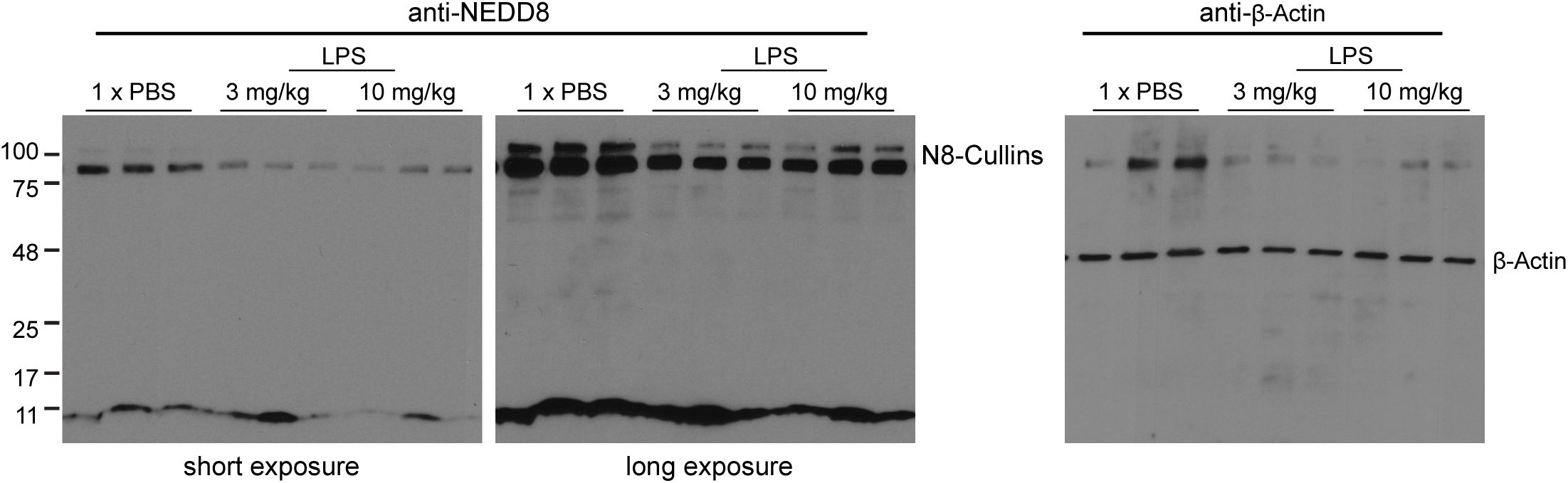
Western blot analysis of neddylation in the livers of mice treated with LPS or PBS.

**Figure S7.**
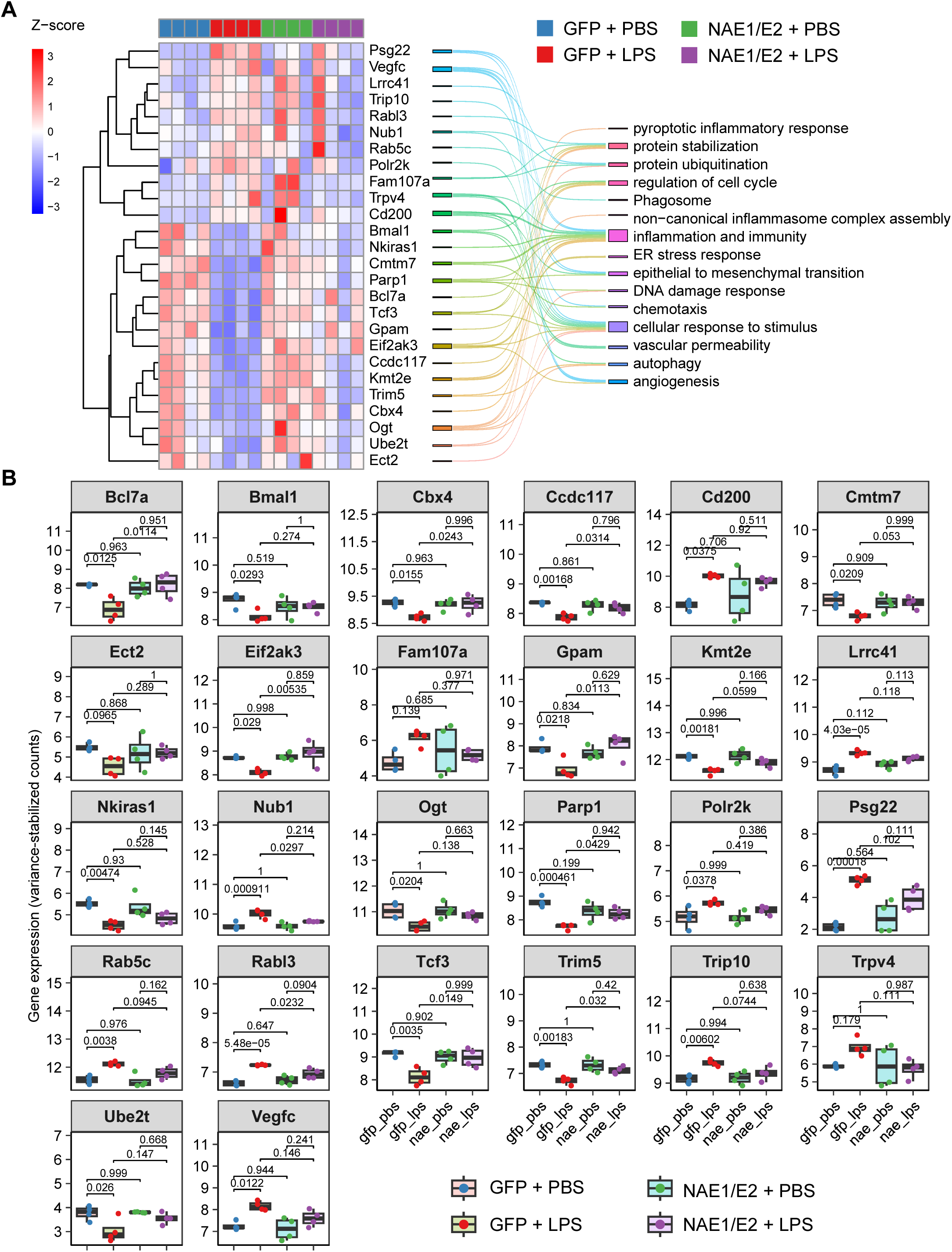
Selected genes that are restored by neddylation E1/E2 expression in the liver ECs of mice treated with LPS. **A**, A heatmap displays the expression levels of 26 selected genes that are differentially expressed in the liver ECs between LPS and PBS in GFP-expressing mice, which were normalized in the liver ECs between LPS and PBS conditions due to the expression of neddylation E1/E2 enzymes. The associated functions of these genes were shown on the right. **B**, Box plots display the expression levels of the 26 selected genes. Statistical significance (*P* < 5.0e-2) was determined by two-way Anova followed by Tukey HSD post-hoc test.

**Figure S8.**
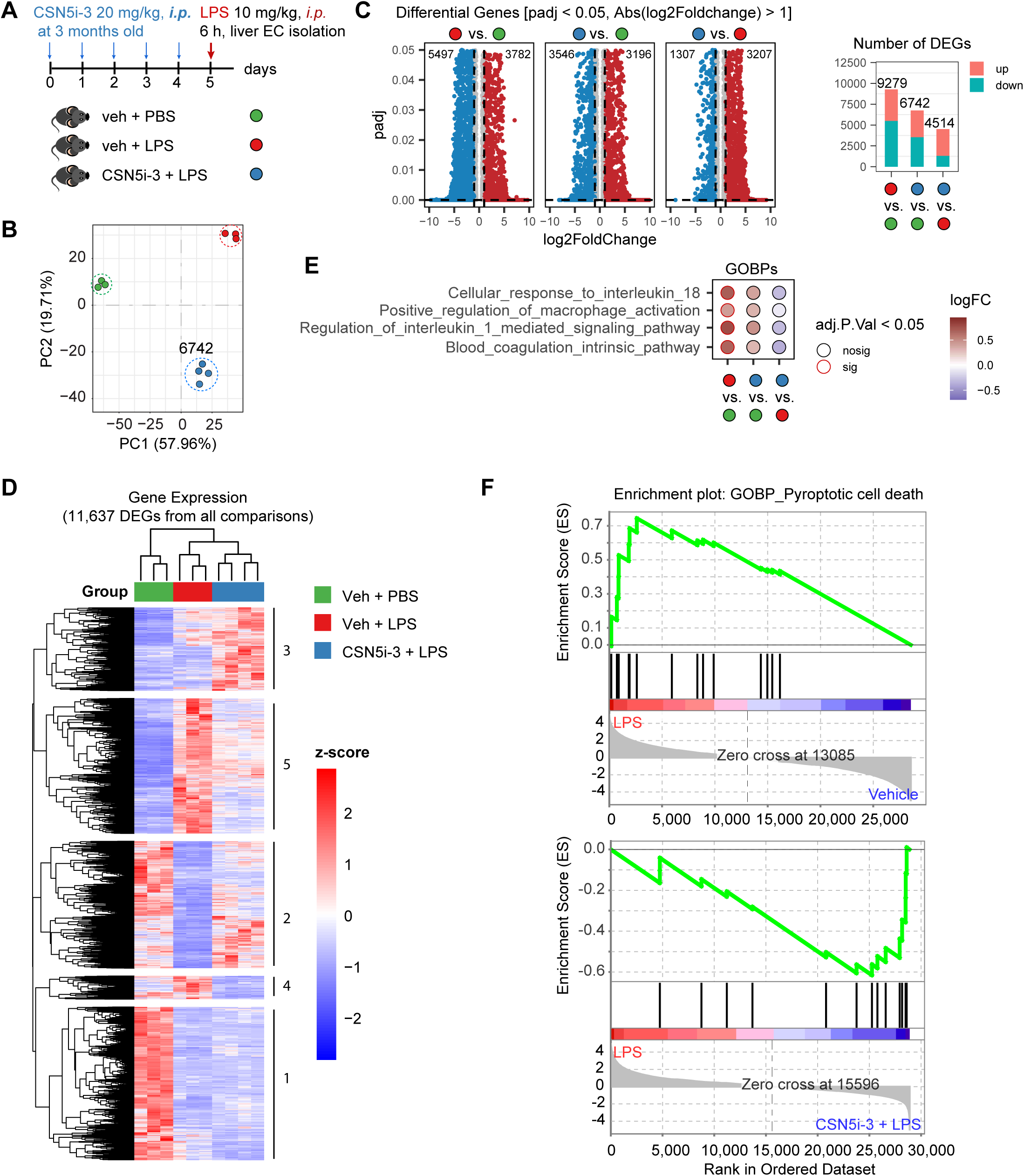
The COP9 signalosome (CSN) inhibitor CSN5i-3 partially reverses LPS-induced endothelial transcriptomic reprogramming. **A**, Schematic representation of the experimental workflow for assessing the effects of CSN5i-3 on LPS-treated endothelial transcriptome. Liver ECs were isolated from 10 mice (n = 3, 3 and 4 mice for vehicle + PBS, vehicle + LPS, and CSN5i-3, respectively). The total RNAs after ribosomal RNAs depletion were used for bulk RNA-seq analysis. **B**, Principal component analysis reveals the global transcriptomic differences among the three groups of mice in A. Each dot represents ECs from one liver. **C**, Volcano plots of differentially expressed genes comparing the indicated groups and a bar plot showing the total number of DEGs in each comparison. **D**, A heat map displays z-scores of normalized expression levels of 11,637 DEGs (*P*adj < 0.05, |log2Foldchnage| > 1) for all comparisons among groups. **E**, Gene Set Variation Analysis identified four up-regulated Gene Ontology Biological Processes (GOBPs) by LPS that are attenuated by CSN5i-3. The color of circle edges reflects adjusted *P* value (*P*adj). Circles are filled with colors that represent the values of log2FC (log2foldchange) of GOBPs for each comparison. **F**, Enrichment plots showing that GOBP pyroptotic cell death is upregulated by LPS and is downregulated by CSN5i-3 in the hepatic endothelium.

### Supplemental Methods

#### Tissue fixation and paraffin embedding

Mice were transcardially perfused with PBS to clear blood from the vasculature prior to tissue collection. The designated liver lobe was dissected and immersion-fixed in 10% neutral buffered formalin (NBF) at 4°C for 24 hours; for a subset of mice, lungs were fixed by intratracheal perfusion with 10% NBF prior to dissection. Following fixation, tissues were rinsed in PBS, dehydrated through a graded ethanol series, cleared in xylene, and infiltrated with paraffin wax using a standard tissue processor. Tissues were embedded in paraffin blocks with consistent orientation and sectioned at 10 μm thickness using a rotary microtome. Sections were mounted on positively charged glass slides and dried before storage at room temperature until staining.

#### Protein Extraction and Western Blot Analysis

Total protein was extracted from HUVECs, mouse liver and lung tissues, or endothelial cells isolated from mouse livers (1 to 2 million cells per mouse liver) using RIPA lysis buffer supplemented with protease and phosphatase inhibitors. Protein concentrations were determined using the Pierce BCA protein assay. Samples were denatured in reducing Laemmli SDS-sample buffer (Boston Bioproducts, <u>#NC9566545</u>) and resolved on SDS-PAGE gels at 120V for 60–70 minutes. Proteins were transferred to nitrocellulose membranes using a semi-dry transfer system. Membranes were blocked with 5% non-fat milk in 1 x Tris-buffered saline with 0.05% Tween-20 (TBST) for 30 minutes at room temperature, followed by incubation with primary antibodies in blocking buffer on rocking shaker for 12-24 h at 4°C (antibodies and dilutions are listed the Major Resources Tables). The membranes were washed in 1 x TBST, and incubated with anti-rabbit (Cell Signaling Technology, 7074) or anti-mouse (Cell Signaling Technology, 7076) HRP-conjugated secondary antibodies in the blocking buffer. After washing, the membranes were incubated with either Pierce ECL Western Blotting Substrate (Thermo Scientific, 32109) or Cytiva Amersham ECL Prime Western Blotting Detection Reagent (Cytiva, RPN2236). X-ray films were used to detect chemiluminescent signals. ImageJ (Fiji) software was used to quantify protein bands. Protein loading controls GAPDH or β-actin were used to normalize the levels of protein of interest.

#### Hematoxylin and Eosin (H&E) Staining

Tissue sections were deparaffinized by immersion in xylene (2x 15 minutes) and then rehydrated through a series of ethanol washes (descending from 100% to 75%). Sections were stained with hematoxylin, rinsed in deionized water, and differentiated in acid alcohol followed by a brief rinse in ammonia water (bluing reagent) to develop nuclear staining. Sections were then counterstained with eosin Y, dehydrated through ascending ethanol series, and mounted for brightfield microscopy.

#### TUNEL/Immunofluorescence Staining and Quantitative Imaging

For both TUNEL and immunofluorescence staining, Formalin-fixed, paraffin-embedded tissue sections were deparaffinized by immersion in xylene (2x 15 minutes) and then rehydrated through a series of ethanol washes (descending from 100% to 75%). Slides were then submerged in a citrate buffer and subjected to heat treatment at 97°C for 8 minutes two times, with cooling to 50°C in between. Sections were permeabilized with a PBS solution containing 0.2% (w/v) gelatin and 0.25% (v/v) Triton X-100 twice for 10 minutes. TUNEL labeling was performed with the Elabscience® One-step TUNEL In Situ Apoptosis Kit (#E-CK-A322). Sections were equilibrated with TdT buffer and then incubated with the TdT Enzyme and Labeling Solution for 60 minutes at 37°C, followed by a PBS wash. For TUNEL staining alone, sections were then incubated with 10 μM Hoechst for 5 minutes, washed again, and mounted for fluorescence microscopy imaging. Immunofluorescence labeling was performed following permeabilization (for immunofluorescence alone) or TUNEL labeling (for dual labeling). Sections were next blocked with 5% BSA in the permeabilization solution for 30 minutes, and then incubated with the primary antibody overnight at 4°C. The following day, sections were washed in PBS and then incubated with secondary antibody and 10 μM Hoechst at 37°C for 60 minutes. Slides were washed in PBS and then washed in 10 mM CuSO4 / 50 mM NH4Cl for 10 minutes. Slides were then mounted for fluorescence microscopy imaging. Sections were imaged using the Nikon Ti2 with uniform settings on the fluorescence filters corresponding to the secondary antibodies or the TUNEL fluorophore, up to three channels. Three to five images per section were quantified using ImageJ to count particle number or total area per image of a specific channel. Average count or area per view was computed for each section.

#### Quantitative PCR analysis

Total RNAs were extracted from mouse tissues after homogenization using TRIzol Reagent (Thermo Fisher, 15596026) according to the manufacturer’s instructions. One microgram of RNA was converted to cDNAs using the High-Capacity cDNA Reverse Transcription Kit (Thermo Fisher, 4368814). Quantitative PCRs (qPCRs) were conducted in the CFX Connect Real-Time System (BioRad) using 2 SYBR Green qPCR Master Mix (Bimake, B21203 or Selleck Chem, B21202). Data was normalized by the delta-delta Ct method. Primer sequences are listed in Major Resources Table.

#### Echocardiography

Transthoracic echocardiography was performed in mice using an MX400 ultra-high-frequency linear-array transducer (30-MHz center frequency) coupled to a VEVO 3100 imaging system (FUJIFILM VisualSonics). Mice were anesthetized with 1-2% isoflurane and maintained at a heart rate of approximately 580 beats/min during imaging. M-mode images were acquired in the parasternal short-axis view. LV morphometric and functional parameters were analyzed offline using Vevo LAB software (FUJIFILM VisualSonics). All measurements were obtained from at least 5 consecutive cardiac cycles and averaged for each animal. EF: left ventricle (LV) ejection fraction. FS:LV fractional shortening. LVPW;d: LV end-diastolic posterior wall thickness. Diameters;d: LV end-diastolic diameters.

#### Endothelial Cell Isolation

Tissue was collected from PBS-perfused mice after euthanasia and kept on ice. Pooled liver samples were minced and then digested in 15 mL DMEM (Gibco <u>#11995-065</u>) containing 1 mg/mL Collagenase Type 2 (Worthington <u>#LS004177</u>) and 1 mg/mL Dispase II (Roche #4942078001) at 37°C with shaking for 40 minutes. Cell slurry wes applied to 100 μm nylon filters washed with equal volume DMEM containing 10% FBS, and then centrifuged at 50xg for 6 min at 4°C. The supernatant was collected into a new tube and centrifuged at 500xg for 5 min at 4°C. The cell pellet was resuspended with bead buffer (500 μl PBS containing 0.5% BSA and 2 mM EDTA), and cells were isolated using one of three methods. Method 1: cell suspension was incubated with sheep anti-rat IgG Dynabeads coated with PECAM-1 antibodies (BD 557355). The bead-bound cells were collected using a magnet and washed three times for snap-freezing and storage. Method 2: CD31 MicroBeads (Miltenyi <u>#130-097-418</u>) were added (50 μl per liver or 30 μl per lung), followed by incubation on a rotating mixer at 4°C for 15 minutes. Samples were then centrifuged at 300xg for 10 min at 4°C and supernatant was discarded. Cells were resuspended in 500 μl bead buffer. Miltenyi MS Columns (#130-042-201) were prepared on the magnetic rack and rinsed with 500 μl bead buffer, and then the cell suspension was applied to column and allowed to filter through and then rinsed three times with 500 μl bead buffer. Finally, 1 mL bead buffer was added to the column and cells were expelled with the provided plunger. An additional 9 mL PBS was added to wash the cells, and then the solution was centrifuged at 300xg for 10 min at 4°C. The cell pellet was washed with 1 mL PBS two additional times, with centrifugation and removal of supernatant in between. Method3: cell suspension from liver was incubated with mouse CD146 microbeads (cat# 130-092-007) from Miltenyi Biotec, and cells were isolated according to the manufacturer’s instructions. Isolated cells were immediately used for single-cell library construction, total RNA isolation for bulk RNA-seq, or stored at - 80°C.

#### Bulk RNA-seq and data analysis

CD31^+^ or CD146^+^ cells were isolated from the livers as described above in the Method2 of endothelial cell isolation. Cells were lysed in TRIzol™ Reagent for total RNA extraction. Total RNAs were sent to Novogene for directional library preparation and pair-end 150 bp sequencing with 12 Gigabases of raw data per sample. Bulk RNA-seq data were analyzed by Novogene following the standard pipeline, including quality control of the raw sequencing reads, alignment of the reads to a reference genome, quantification of gene expression levels, and differential gene expression analysis to find genes that change between samples. Ingenuity Pathway Analysis was performed by the Bioinformatics and Systems Biology Core at the University of Nebraska Medical Center.

Total RNAs of livers from GSDM silencing experiment were sent to GENEWIZ for library construction and sequencing. Libraries were prepared by using the QIAGEN FastSelect rRNA HMR Kit (Qiagen, Hilden, Germany) and the NEBNext Ultra II RNA Library Preparation Kit for Illumina, following the manufacturer’s recommendations (NEB, Ipswich, MA, USA). Raw FASTQ files were processed to remove adapter sequences using trim-galore (0.6.10). Trimmed FASTQ files were aligned to the mouse reference genome mm39 (GRCm39) using hisat2 (2.2.1). FeatureCounts (2.0.3) was used to count the read numbers mapped to each gene. Differential expression analysis was performed using the DESeq2 (1.46.0) package. PCA plot was generated using the PCAtools package (2.18.0). Volcano plots and dot plots were generated using ggplot (4.0.1), and the heatmap was generated using pheatmap (1.0.13) in R (4.4.3) and Rstudio 2025.09.2+418.

#### Gene Set Enrichment Analysis

To perform Gene Set Enrichment Analysis for snRNA-seq or scRNA-seq data, cells in a cluster were randomly assigned to three replicates per genotype. Gene expression counts of all cells in a replicate were aggregated for each gene using the AggregateExpression function in Seurat to create the gene expression matrix of the six pseudo-bulk samples. The normalized gene counts were used for Gene Set Enrichment Analysis using the GSEA v4.4.0 software for Windows. Alternatively, differentially expressed genes were input into Enrichr^3^ to perform functional enrichment analysis. To perform Gene Set Enrichment Analysis for bulk RNA-seq data, DESeq2-normalized counts were used as the input and analyzed using the default parameters with gene set permutation type for the Gene Set Enrichment Analysis using the GSEA v4.4.0 software for Windows. The gene set ‘pyroptosis’ was manually curated and contains the genes listed in the Supplementary Table 1. The gene set “vascular EC core gene set” was generated previously^4^. The ggplot2 package was used to produce plots for data visualization in R (version 4.3.3) and Rstudio (2023.12.1 Build 402).

#### TMT-Labeling Proteomics

HUVECs cultured in 100 mm plate were treated with DMSO or the neddylation inhibitor MLN4924 at 100 nM for 24 hours. Cells were detached with trypsin, washed five times with ice-cold 1 x dPBS, and snap-frozen on dry ice. The samples were submitted to the Proteomics and Metabolomics Facility at our institute and processed for labelling with TMT10plex™ Isobaric Label Reagents and quantitative proteomics as described in our previous studies^2^.

Samples were lysed in RIPA buffer supplemented with 5 mM DTT, Complete EDTA-free protease inhibitor cocktail (Roche), and 2× PhosSTOP phosphatase inhibitors (Roche). Lysates were incubated on ice for 20 min, followed by heating at 95°C for 5 min with shaking in a thermomixer. Following centrifugation, supernatants were transferred to fresh tubes and protein concentration was determined using the CBX Protein Assay Kit (G-Biosciences).

For each sample, 130 μg of protein was alkylated with 20 mM iodoacetamide for 40 min in the dark and excess reagent was quenched with DTT. Proteins were precipitated overnight with acetone and the resulting pellets were washed three times with 70% ethanol. Protein pellets were resuspended in 1% sodium deoxycholate, 100 mM EPPS buffer (pH 8.5), and digested sequentially with Lys-C (1:100 enzyme-to-substrate ratio) for 3 h and trypsin (1:100) overnight at 37°C.

Peptide digests were acidified with 0.5% trifluoroacetic acid (TFA), incubated for 30 min, and centrifuged. Supernatants were transferred to fresh tubes, dried, and resuspended in 100 μL of 100 mM EPPS (pH 8.5). Peptides were labeled using TMT 10-plex reagents (Thermo Fisher Scientific), with 130 μg of peptide labeled at a 1:2 peptide-to-reagent ratio. Following labeling, samples were pooled, acidified to 1% formic acid, and desalted using 50 mg Sep-Pak C18 solid-phase extraction cartridges (Waters Corporation).

For offline fractionation, 200 μg of pooled peptide material was separated into 96 fractions using high-pH reverse-phase chromatography on an ACQUITY UPLC BEH C18 column (1.7 μm, 2.1 × 100 mm; Waters Corporation) operating at pH 10. Fractions were concatenated into 12 final fractions prior to LC-MS/MS analysis as previously described (Yang et al., 2012; 10.1016/j.jprot.2023.104839).

Each fraction was analyzed by nanoLC-MS/MS using an RSLCnano system (Thermo Fisher Scientific) coupled to an Orbitrap Eclipse Tribrid mass spectrometer (Thermo Fisher Scientific) as previously described (Olanrewaju et al, 2023, https://doi.org/10.3389/fpls.2023.1260429). Samples were initially loaded onto an Acclaim PepMap 100 trap column (75 μm × 2 cm) for 3 min at 5 μL/min and subsequently separated on an analytical column. Mobile phase A consisted of water containing 0.1% formic acid, while mobile phase B consisted of acetonitrile containing 0.1% formic acid.

Chromatography was performed on a longer Peptide CSH C18 column (130 Å, 1.7 μm, 75 μm × 500 mm; Waters Corporation) at 250 nL/min with a linear gradient of 5% to 30% solvent B over 130 min. MS1 spectra were acquired in the Orbitrap, and MS2 spectra were generated using higher-energy collisional dissociation (HCD) at 32% normalized collision energy. RTS settings were similar to those used in 2022, except that phosphorylation was excluded from the variable modifications. FAIMS compensation voltages were set at −45, −60, and −75 V.

Raw data were processed in Proteome Discoverer v2.4 (Thermo Fisher Scientific). Spectra were searched against the UniProt human database (release 20210508; 77,027 entries) using the Sequest HT search engine with trypsin specified as the digestion enzyme. Search parameters included a precursor mass tolerance of 10 ppm and a fragment mass tolerance of 0.66 Da. Methionine oxidation and phosphorylation of serine, threonine, and tyrosine residues were specified as variable modifications, while carbamidomethylation of cysteine and TMT labeling of lysine residues and peptide N-termini were specified as fixed modifications. Peptide-spectrum matches were validated using Percolator with a posterior error probability threshold of 0.01, and a target-decoy approach was used to control the false discovery rate at 1%.

Protein quantification was performed in Proteome Discoverer using SPS-MS3 reporter ion intensities with a co-isolation threshold of 50%, an average signal-to-noise ratio threshold of 10, and an SPS mass match threshold of 65%. Reporter ion abundances were normalized based on total peptide loading across samples. Protein abundances were calculated from summed peptide abundances for each biological replicate, and protein ratios were determined using the geometric median of replicate ratios. Differential protein abundance was assessed by analysis of variance (ANOVA) and resulting p-values were adjusted for multiple testing using the Benjamini-Hochberg false discovery rate correction.

#### FITC-dextran permeability assay

FITC-dextran (70 kDa; Sigma-Aldrich, cat# 46945) was dissolved in saline at 50 mg/ml. It was intravenously administered in mice (10 mg/mouse)^1,2^. FITC-dextran was allowed to circulate for 20 min. Mice were euthanized to collect for fresh liver OCT embedding and cryo-sectioning. After anti-CD31 staining, four images of random fields of view per mouse were acquired for quantification using ImageJ.

#### Nuclei isolation from snap-frozen livers

The isolation of nuclei from snap-frozen livers was performed as previously described ^3^. Briefly, about 200 mg of liver tissue was pooled from the same anatomic lobe of five mice for each nucleus isolation. Allow the tissue to thaw for 5 minutes in a petri dish over ice containing ice-cold lysis buffer. Mince the tissue into 2-4 mm pieces, add 2 ml of ice-cold PBS, and mix gently by pipetting up and down 10 times. The lysates were filtered through a 30 µm smart strainer for spin to collect the pellet, which was washed for nuclei isolation. Ten microliters of the isolated nuclei were stained with Hoechst 33342 for nucleus counting and quality assessment. The yield of nuclei is about 5,000 per mg of liver tissue. The nuclei were diluted with the wash and resuspension buffer to achieve a concentration of 1,000 nuclei per microliter.

#### Single-cell and single-nucleus RNA-seq and data analysis

Sequencing libraries were prepared using the 10x Genomics Chromium Single Cell 3’ Reagent Kit at our Flow Cytometry and Single-Cell Genomics Core Facilities and sequenced on an Illumina Novaseq 6000 instrument at the Genome Sequencing Facility at the Kansas University Medical Center. Raw data were processed using Cell Ranger and subsequent analyses were performed using Seurat v.5.3.0. In brief, FASTQ files containing sequenced reads were mapped to the mouse reference genome (mm10). After Seurat objects were generated, the SoupX package (1.6.2) was used to remove ambient RNA; low quality cells were removed in the merged Seurat object after quality control filtering [500 < nCount_RNA < 50000, 200 < nFeature_RNA < 7500, reads mapped to hemoglobin genes < 3%, and mitochondrial reads < 10% (for scRNA-seq); nCount_RNA > 500, nFeature_RNA > 200, reads mapped to hemoglobin genes < 3%, and mitochondrial reads < 5% (for snRNA-seq)]; and the scDblFinder package (1.20.2) was used to detect doublets. After gene expression counts were normalized and scaled, the Principal Component Analysis was performed to reduce data dimensionality, and the batch effects were corrected using Harmony (1.2.3). The Harmony-corrected data were analyzed for graph-based clustering using the top 20 principal components at a resolution of 0.1 (scRNA-seq) or 0.2 (snRNA-seq). Cell clusters were manually annotated based on marker genes of each cell type from published single-cell transcriptome studies^4–6^. To identify differentially expressed genes between two phenotypes, we performed two different methods. Method1: The cells of the EC cluster in each sample were randomly split into three replicates, and gene expression counts were then aggregated to generate pseudo-bulk data for each sample. Aggregated gene expression counts were used as input data for DEseq2 (1.42.1) analysis. Method2: Seurat package FindMarkers function was used to identify genes that are different between two phenotypes. By default, Seurat performs differential expression testing based on the non-parametric Wilcoxon rank sum test.

#### Endothelial subclustering and Cell trajectory analysis

Briefly, three functions—SelectIntegrationFeatures, FindIntegrationAnchors, IntegrateData—were used to integrate the endothelial cell populations of scRNA-seq and snRNA-seq for subclustering. The Seurat object was then processed using the standard procedures involving scaling expression data, running linear and non-linear dimensional reduction, building a shared nearest neighbor graph, and partitioning cells into distinct phenotypic clusters. The top 30 components were used to construct a low-dimensional UMAP embedding and identify cell clusters at the resolution of 0.3.

Cell trajectory and pseudotime analyses were performed using the Slingshot package (v2.14.0)^7^. The Seurat object was converted into a SingleCellExperiment object. Slingshot was then used to infer cell lineages by employing a clustering-derived minimum spanning tree to outline lineage structures. Smooth principal curves were simultaneously fit to these lineages to define the paths, with the starting root cluster identified by default. A continuous pseudotime value was assigned to each individual cell. The subclusters 5, 6, and 7 were not used for cell trajectory analysis because there are only 14, 21, and 15 cells for the iECKO condition. The cell death and inflammation pathway activities were calculated using the AddModuleScore function in the Seurat R package. It calculated the average expression levels of a specific set of genes involved in cell death or inflammation in individual cells. The activities of both pathways in all cells were plotted as the function of pseudotime.

#### Cell‒cell communication analysis

The R package CellChat (v2.2.0)^8,9^ was used to infer cell-cell interactions within the snRNA-seq dataset. Analysis followed the standard pipeline using default parameters and the complete `CellChatDB.mouse` database. To compare the *Nae1* fl/fl and iECKO conditions, separate CellChat objects were constructed for each condition, merged into a single list, and analyzed using quantitative comparative functions to evaluate shifts in signaling pathways. Specifically, `netAnalysis_computeCentrality` was used to contrast global patterns and specific pathways, `computeCommunProbPathway` predicted major incoming and outgoing signals, and `netAnalysis_signalingRole_heatmap` visualized signaling strength across cell types. Finally, interaction counts, differential networks, and changes in signaling pathways were identified and visualized.

#### AAV9 Vector Design, Production, and *In Vivo* Delivery

Adeno-associated virus, serotype 9 (AAV9) constructs expressing neddylation E1 enzymes (NAE1 and UBA3) and E2 enzymes (UBE2M and UBE2F) were generated as follows. First, the control plasmid pAAV-ICAM2-GFP was generated by excising the TBG promoter from pAAV.TBG.PI.eGFP.WPRE.bGH (Addgene, 105535) with restriction enzymes AscI and NotI and replacing with a synthesized ICAM2 promoter. Next, 3xFLAG-UBE2M and HA-UBE2F were cloned into the pAAV-ICAM2-GFP vector using NotI/HindIII and NotI/AgeI restriction sites, respectively. Additionally, HA-NAE1 and FLAG-UBA3 were cloned from mouse cDNA using synthesized primers and then inserted into the pAAV-ICAM2-GFP vector. Specifically, mouse *Nae1* cDNA were amplified using primers forward-NotI-SalI: agtGCGGCCGCGGTGGTGTCGACATGGCGCAGCCAGGGAAGATAC

and reverse-BamHI: agtGGATCCCTACAACTGGAAAGTTGCAGAAG. *Nae1* gene was inserted into pAAV-ICAM2-GFP between NotI and BamHI sites to replace GFP. Then the HA-tag adaptor was annealed from two oligos HA_s: 5’ ggccgc GCCACCATGtacccatacgatgttccagattacgctGGATCCGGCGGGTCCg

HA-as: 5’ tcgacGGACCCGCCGGATCCagcgtaatctggaacatcgtatgggtaCATGGTGGCgc

and inserted between NotI and SalI sites in pAAV-ICAM2-NAE1, resulting in pAAV-ICAM2-haNAE1. Mouse *UBA3* cDNA was amplified using primers mUBA3-NotI_SalI: agtGCGGCCGCGGTGGTGTCGACATGGCGGATGGCGAGGAGC

mUBA3-HindIII: aca aagcttTTAAGTAAAATGAAGTTTGAATAG.

*UBA3* gene was inserted into pAAV-ICAM2-GFP between NotI and HindIII sites to replace GFP. Then the Flag-tag adaptor was annealed from two oligos Flag_s: 5’ ggccgcGCCACCATGGACTACAAGGATGACGACGATAAGGGATCCGGGTCGGGCTCCg

Flag-as: 5’ tcgacGGAGCCCGACCCGGATCCCTTATCGTCGTCATCCTTGTAGTCCATGGTGGCgc and inserted between NotI and SalI sites in pAAV-ICAM2-UBA3, resulting in pAAV-ICAM2-flagUBA3.

The E1 plasmids pAAV-ICAM2-haNAE1 and pAAV-ICAM2-flagUBA3 were mixed in a 1:1 ratio, and the E2 plasmids pAAV-ICAM2-flagUBE2M and pAAV-ICAM2-haUBE2F were likewise mixed 1:1. The E1 and E2 constructs were used to generate AAV9 as described below. All sequences are listed in the Major Resources Tables.

For AAV9-mediated shRNA delivery, AAV-H1-shRNA also known as scAAV-RSV-GFP-H1^10,11^ from Dr. Ira Tabas’s laboratory (produced by Dr. Mark Kay’s lab) was used as the vector to express non-targeting, GSDMD, and GSDME shRNAs. It has been used to effectively knockdown gene expression in murine liver^12,13^. Oligonucleotides were designed, and the forward and reverse oligos were annealed and ligated into the vector digested with BbsI. These plasmids were called pAAV-shNT, pAAV-shGSDMD and pAAV-shGSDME. The different shRNA plasmids were co-transfected into HepG2 cells with GSDMD and GSDME overexpression vectors to test effectivity at knocking down GSDMD or GSDME, and the most effective shRNA(s) for each target were selected for virus production; the two GSDMD shRNAs were mixed in a 1:1 ratio. The pAAV-shNT, pAAV-shGSDMD, and pAAV-shGSDME plasmids were used to generate AAV9 as described below. All sequences are listed in the Major Resources Tables.

To produce recombinant AAV9, the AAVpro 293T cell line (Takara, 632273) is used. Briefly, AAVpro 293T cells were cultured in twenty T-175 flasks. Cells at 80% confluency were co-transfected with an equimolar ratio of shRNA- or exogenous gene-expressing plasmid (20 μg per flask), pAAV2/9n (Addgene, 112864), and pHelper vector from the AAVpro Helper Free System (Takara, 6673). Polyethylenimine (PEI) (Polysciences, 23966-1) was used for transfection with a 3:1 ratio of PEI to plasmid DNA (w/w). After 24-hour transfection, media was replaced with DMEM (Gibco, # 11965-092) supplemented with 2% FBS, 1% penicillin-streptomycin (Gibco, # 15140-122), and 1% L-glutamine (Gibco, # 25030-081). After 48 h, The supernatant was collected, cleared of cellular debris, and stored at 4°C, while fresh medium was added to the flasks. A further 48 h later, cells were collected in the supernatant and centrifuged at 3000 x g for 20 min to pellet the cells. The supernatant was saved, and the cell pellets were resuspended in 20 ml AAV freezing buffer (50 mM Tris-HCl pH 7.4, 150 mM sodium chloride, 2 mM magnesium chloride). After three freeze-thaw cycles, cell lysates were clarified by centrifugation at 2500 g for 30 min. Clarified cell lysates and saved supernatants media were combined, mixed with 25% volume PEG extraction solution [40 % (w/v) PEG-8000, 2.5 M NaCl in sterile water], and incubated on ice for 2 h. The mixture was spun at 2500 x g for 30 min at 4°C. The pellets were dissolved in 6.5 ml AAV freezing medium, treated with 50 units per ml of TurboNuclease (Eton Bioscience, 1400010050) at 37°C for 1 h. Cesium chloride (CsCl) gradient centrifugation was performed to purify the virus as previously described^14^ with minor modifications. Briefly, the nuclease-treated mixture (6.5 ml) was overlaid onto a CsCl solution prepared by dispensing 2.5 ml 1.5 g/ml CsCl below 2.5 ml 1.3 g/ml CsCl in a standard ultraclear tube (Beckman, 344059) and centrifuged at 29,028 RPM (104,000 g) in an SW41 Ti rotor. After centrifugation, fractions containing virus particles were collected and dialyzed in Slide-A-Lyzer Dialysis Cassettes (Thermo Scientific, 66330) at 4°C for at least 12 h with three buffer (1 dPBS) exchanges. The virus solution was then concentrated with an Amicon Ultra-4 Centrifugal Filter Unit (Millipore, UFC810024). Viral titer was determined by qPCR according to the protocol “AAV Titration by qPCR Using SYBR Green Technology” from Addgene.

To express shRNA in mice, NAE flox mice (Cre- and Cre+) were intravenously injected with AAV9-shRNA-NT, or a mixture of AAV9-shGSDMD and AAV9-shGSDME via tail vein. After 4 weeks, tamoxifen was administered (50 mg/kg for 5 consecutive days) to induce NAE iECKO. After a further 24 days post-TAM, mice were euthanized for blood and tissue collection.

For expression of neddylation E1/E2 genes, wild type C57BL/J6 mice were intravenously injected with AAV9-ICAM2-GFP or a mixture of AAV9-ICAM2-E1 and AAV9-ICAM2-E2. After 4 weeks, mice were injected intraperitoneally with LPS or PBS at 10 mg/kg, then euthanized 6 h later for blood and tissue collection.

#### Gene Set Variation Analysis

Bioconductor package Gene Set Variation Analysis (GSVA 3.22) was used. The mouse Gene Ontology (GO) Biological Process (BP) gene sets (m5.go.bp.v2025.1.Mm.symbols.gmt) were downloaded from the Molecular Signatures Database (MSigDB). We calculated GSVA enrichment scores for these gene sets using the normalized and log1p transformed gene counts of each replicate. We then performed a differential expression analysis at the gene set level using the Bioconductor package limma (3.62.2). The higher absolute mean differences (log2FoldChange) of GSVA scores between groups and a low adjusted p-value identify robust biologically meaningful pathways that are upregulated (positive values) or downregulated (negative values).

#### Public GEO datasets analysis

We obtained and analyzed four bulk RNA-seq datasets from the NCBI Gene Expression Omnibus (GEO) database. The datasets contain gene expression in human carotid arteries with atheroma plaque or distant macroscopically intact tissue (GSE43292), with early or advanced atherosclerotic plaque (GSE28829), carotid arteries with or without atherosclerotic lesions (GSE100927), whole blood from healthy or COVID positive individuals (GSE217948). Normalized counts in TPM were plotted for GSE217948. The datasets GSE100927, GSE43292, and GSE28829 were analyzed using GEO2R after the groups were defined. The expression values of genes were copied into a CSV file. Because the expression values were log2 transformed, we performed an inverse log2 transformation in R to determine the original normalized intensity signal for each gene. Subsequently, the normalized intensity signal values were plotted using the ggviolin function from the ggpubr (0.6.3) package in R (4.4.3).

To correlate the expression of NAE1, CUL1, and CUL3 with other genes, we downloaded and analyzed two scRNA-seq datasets of human arteries, including GSE131778 and GSE198750. Processed data files were used to create Seurat objects using Seurat package (5.3.0). The low-quality cells were removed in the merged Seurat object after quality control filtering [500 < nCount_RNA < 50000, 200 < nFeature_RNA < 6000 (5000 for GSE198750), reads mapped to hemoglobin genes < 3%, and mitochondrial reads < 10%]. After gene expression counts were normalized and scaled, the Principal Component Analysis was performed to reduce data dimensionality, and the scDblFinder package (1.20.2) was used to detect doublets. The batch effects were corrected using Harmony (1.2.3). The Harmony-corrected data were analyzed for graph-based clustering using the top 20 principal components at a resolution of 0.1. Cells were annotated as ECs based on the expression of marker genes, including PECAM1, CDH5, and vWF. ECs from each dataset were merged to create a new Seurat object. The expression of each gene was calculated using the AverageExpression function in Seurat grouped by each sequencing sample (donor) from the “data” layer. “Zero” values were replaced with “NA” before the Spearman’s correlation coefficients along with their corresponding asymptotic p-values were computed using the rcorr function from the Hmisc package (5.2-5). The correlation heatmap was generated using the pheatmap (1.0.13) in R (4.4.3) and Rstudio 2025.09.2+418.

#### LPS Intraperitoneal Injection

Male C57BL/6J mice (Jackson Laboratory) were injected intraperitoneally with lipopolysaccharide (LPS; AdipoGen, <u>#IAX-100-012-C500</u>; *E. coli* O111:B4) dissolved in dPBS at 0.3 or 1 mg/mL and administered at 10 μl/g body weight for a total dose of 3 or 10 mg/kg, respectively, or an equal volume of dPBS vehicle as a control. At six hours post-injection, mice were euthanized and CD31+ liver endothelial cells were isolated from fresh livers as described below (see Endothelial Cell Isolation) for downstream analysis.

### Major Resources Tables

#### Mouse Models Used in This Study

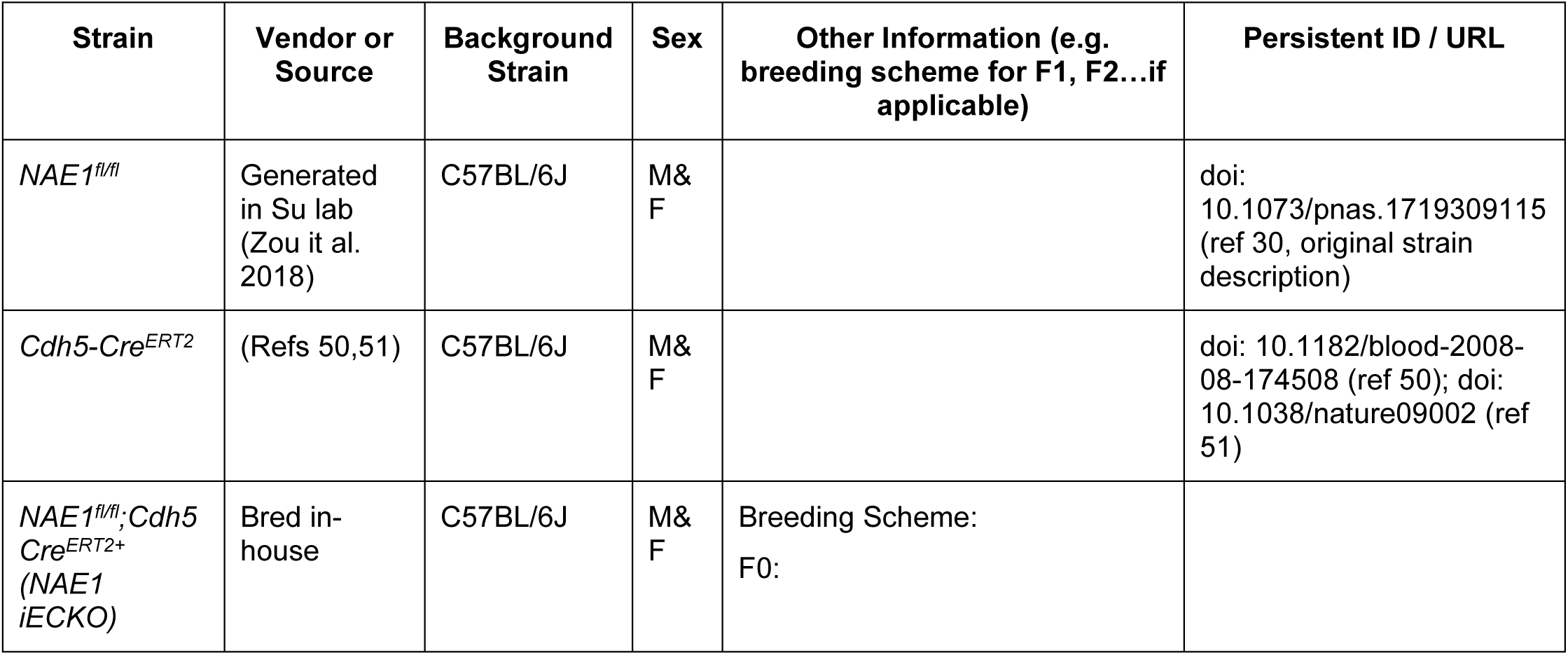
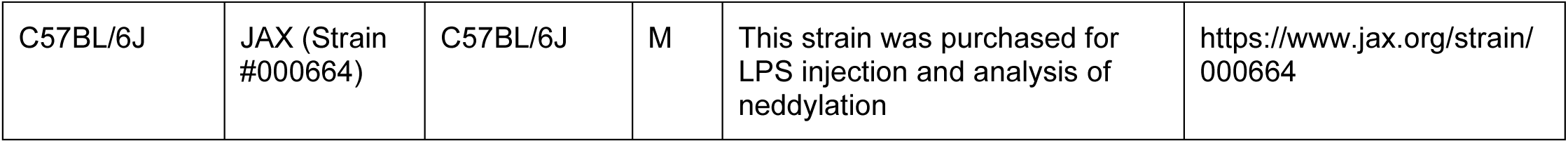

| Strain | Vendor or Source | Background Strain | Sex | Other Information (e.g. breeding scheme for F1, F2...if applicable) | Persistent ID / URL |
| --- | --- | --- | --- | --- | --- |
| *NAE1<sup>fl/fl</sup>* | Generated in Su lab (Zou et al. 2018) | C57BL/6J | M&F |  | doi: 10.1073/pnas.1719309115 (ref 30, original strain description) |
| *Cdh5-Cre<sup>ERT2</sup>* | (Refs 50,51) | C57BL/6J | M&F |  | doi: 10.1182/blood-2008-08-174508 (ref 50); doi: 10.1038/nature09002 (ref 51) |
| *NAE1<sup>fl/fl</sup>;Cdh5-Cre<sup>ERT2+</sup>* (NAE1 iECKO) | Bred in-house | C57BL/6J | M&F | Breeding Scheme: F0: |  |
| C57BL/6J | JAX (Strain #000664) | C57BL/6J | M | This strain was purchased for LPS injection and analysis of neddylation | https://www.jax.org/strain/000664 |

#### Antibodies

| Target antigen | Vendor or Source | Catalog # | Working concentration | Lot # (preferred but not required) | Persistent ID / URL |
| --- | --- | --- | --- | --- | --- |
| NAE1 | Cell Signaling Technology | 14321S | 1:1000 |  |  |
| NEDD8 | Cell Signaling Technology | 2754S | 1:1000 |  |  |
| Cullin-3 | Bethyl | A301-109A | 1:1000 |  |  |
| HepPar1 | Novus | NBP2-45272 | 1:100 |  |  |
| Mac2 | Cedar Lake | CL8942AP | 1:1000 |  |  |
| CD31 | Dianova | DIA-310 | 1:100 |  |  |
| CD68 | BioLegend | 137001 | 1:500 |  |  |
| Mpo | Abcam | Ab-9535 | 1:25 |  |  |
| VCAM-1 | Abcam | Ab134047 | 1:1000 |  |  |
| E-Selectin | Santa Cruz | Sc-137054 | 1:250 |  |  |
| E-Selectin | R&D | MAB5751 | 1:500 |  |  |
| VE-cadherin | R&D | AF1002-SP | 1:1000 |  |  |
| Claudin-5 | Invitrogen | 34-1600 | 1:200 |  |  |
| β-actin | Cell Signaling Technology | 4970S | 1:5000-1:10,000 | | |
| TER-119 | BioLegend | 116202 | 1:500 |  |  |
| CD41 | Proteintech | 24552-1-AP | 1:800 |  |  |
| NLRP3 | Adipogen | AG-20B-0014 | 1:1000 |  |  |
| GSDMD | CST | 69469 | 1:1000-1:2000 |  |  |
| GSDME | Abcam | Ab215191 | 1:500-1:1000 |  |  |
| Caspase 1 | Adipogen | AG-20B-0042 | 1:1000 |  |  |
| UBA3 | Abcam | Ab124728 | 1:1000 |  |  |
| PECAM-1 (CD31) | BD Pharmingen | 557355 | 40 µl per 200 µl beads |  |  |
| CD31 MicroBeads | Miltenyi Biotec | 130-097-418 | 50 µl/liver, 30 µl/lung |  |  |
| CD146 MicroBeads | Miltenyi Biotec | 130-092-007 | 50 µl/liver |  |  |
| Anti-rabbit IgG, HRP-linked | Cell Signaling Technology | 7074 | 1:5000 |  |  |
| Anti-mouse IgG, HRP-linked | Cell Signaling Technology | 7076 | 1:5000 |  |  |
| Anti-goat IgG, HRP-linked | Invitrogen | 31402 | 1:10,000 |  |  |
| Anti-rat, AF488 | Jackson ImmunoResearch | 712-545-153 | 1:200 |  |  |
| Anti-rat, AF647 | Jackson ImmunoResearch | 112-605-167 | 1:200 |  |  |
| Anti-rabbit, AF488 | Jackson ImmunoResearch | 111-545-144 | 1:200 |  |  |
| Anti-rabbit, AF594 | Jackson ImmunoResearch | 111-585-144 | 1:200 |  |  |
| Anti-mouse, AF488 | Jackson ImmunoResearch | 715-454-151 | 1:200 |  |  |

#### DNA/cDNA Clones

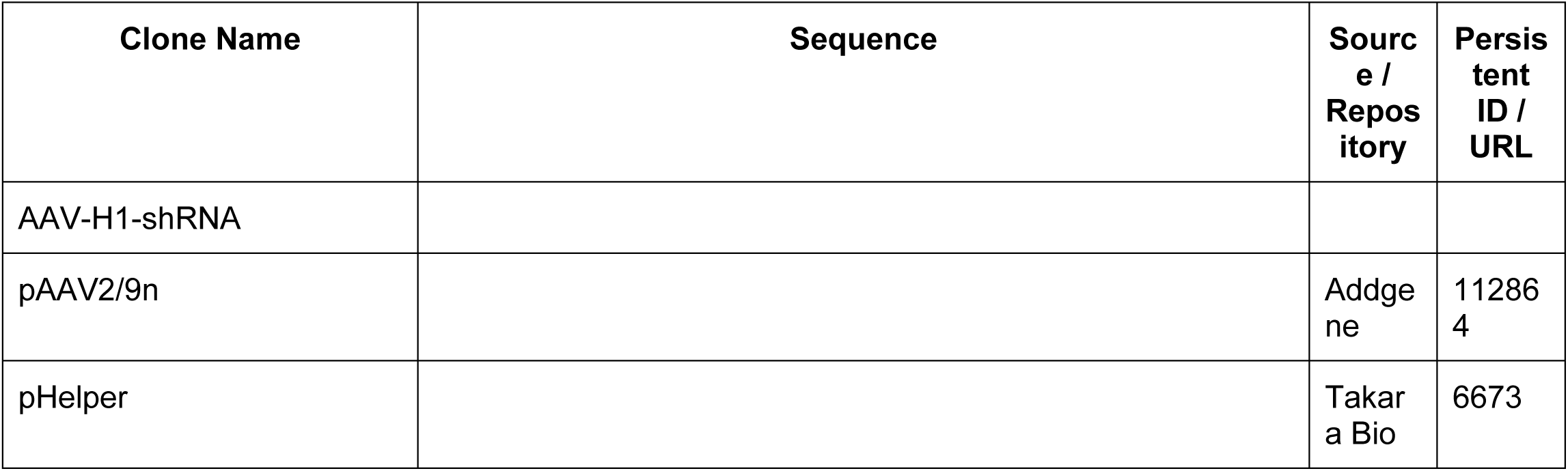
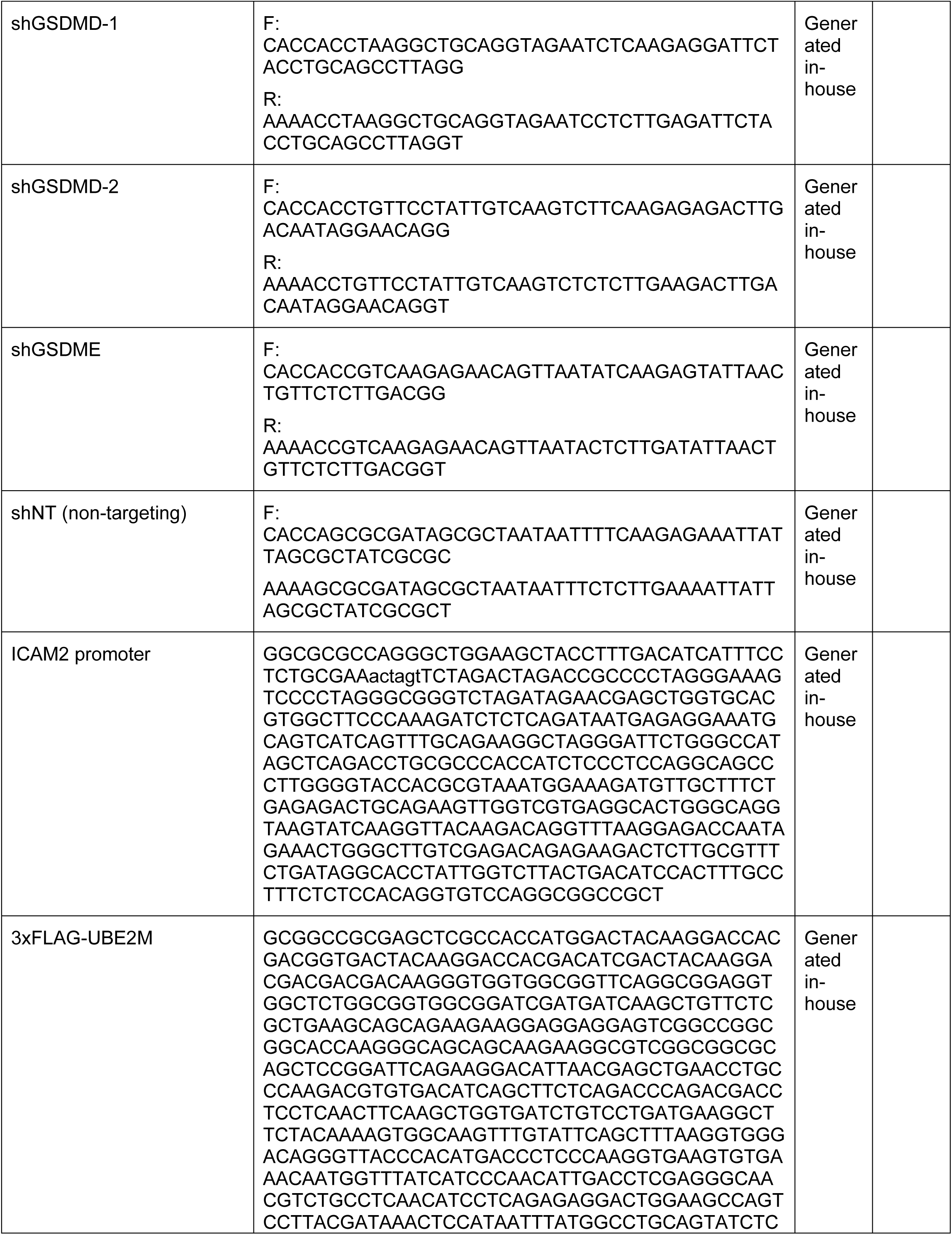
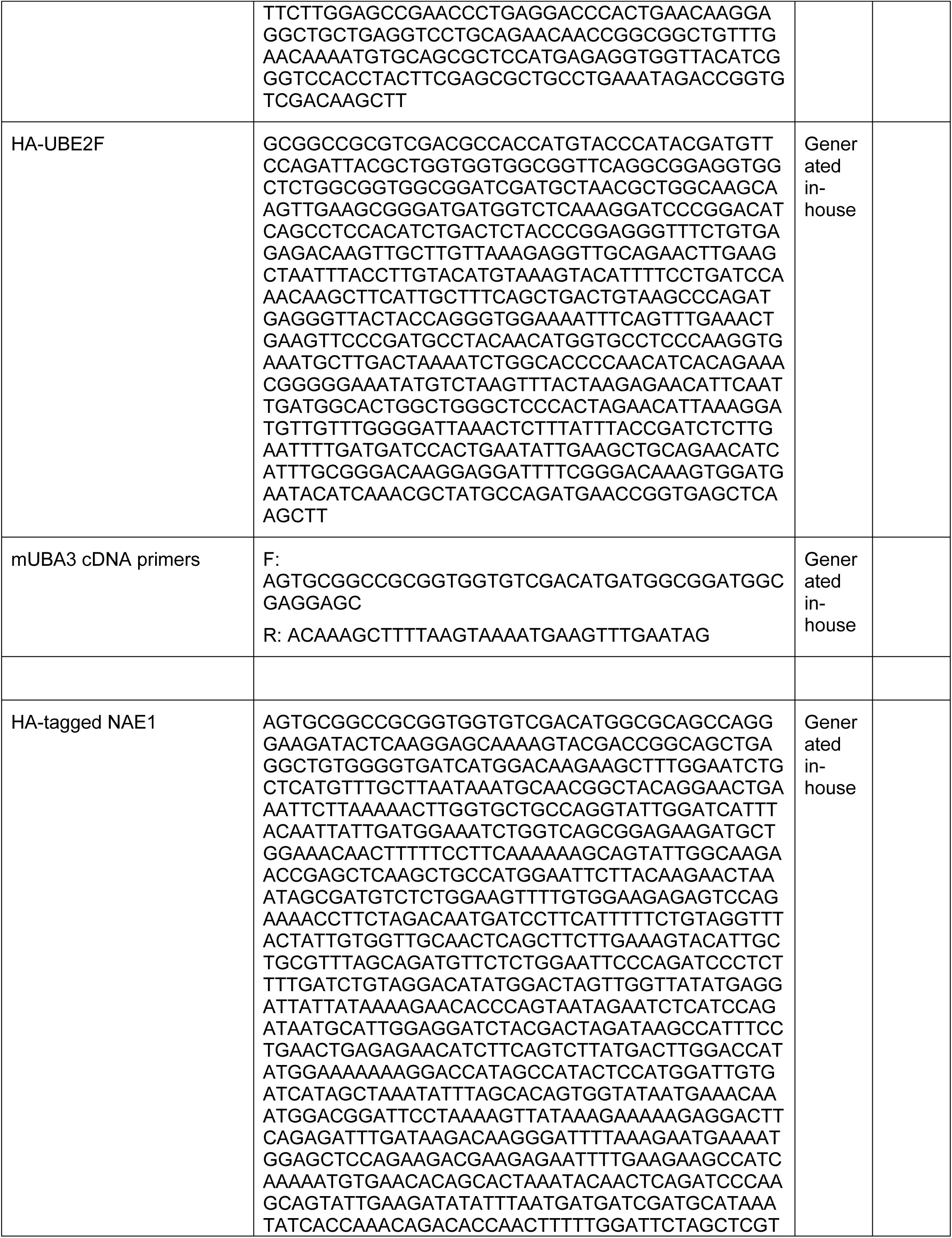
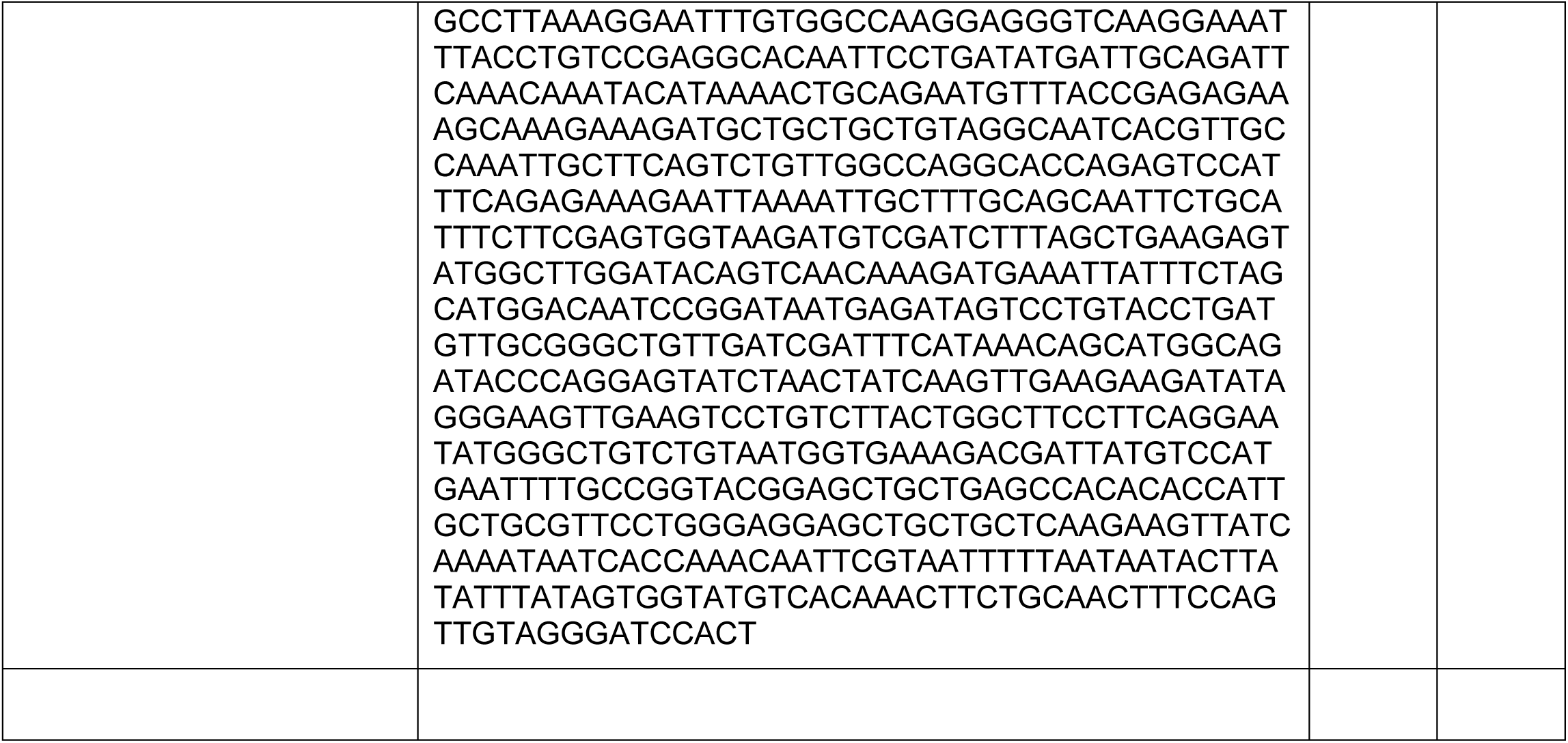

#### Cultured Cells

| Name | Vendor or Source | Sex (F, M, or unknown) | Persistent ID / URL |
| --- | --- | --- | --- |
| HUVEC | Lonza | Unknown | C2519A |
| AAVpro 293T | Takara Bio | Unknown | 632273 |
| HepG2 | ATCC | Male | HB-8065 |

#### Data & Code Availability

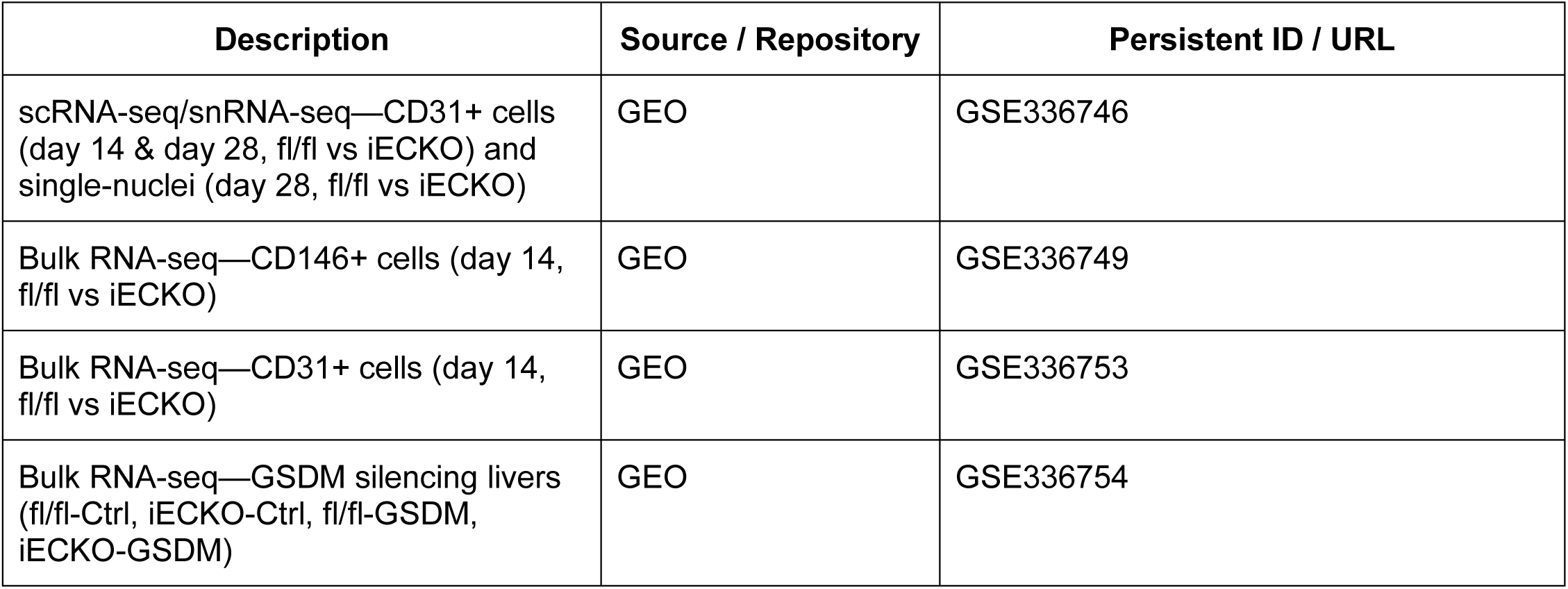

| Description | Source / Repository | Persistent ID / URL |
| --- | --- | --- |
| scRNA-seq/snRNA-seq—CD31+ cells (day 14 & day 28, fl/fl vs iECKO) and single-nuclei (day 28, fl/fl vs iECKO) | GEO | GSE336746 |
| Bulk RNA-seq—CD146+ cells (day 14, fl/fl vs iECKO) | GEO | GSE336749 |
| Bulk RNA-seq—CD31+ cells (day 14, fl/fl vs iECKO) | GEO | GSE336753 |
| Bulk RNA-seq—GSDM silencing livers (fl/fl-Ctrl, iECKO-Ctrl, fl/fl-GSDM, iECKO-GSDM) | GEO | GSE336754 |

#### Chemicals, Peptides, Recombinant Proteins, and Kits

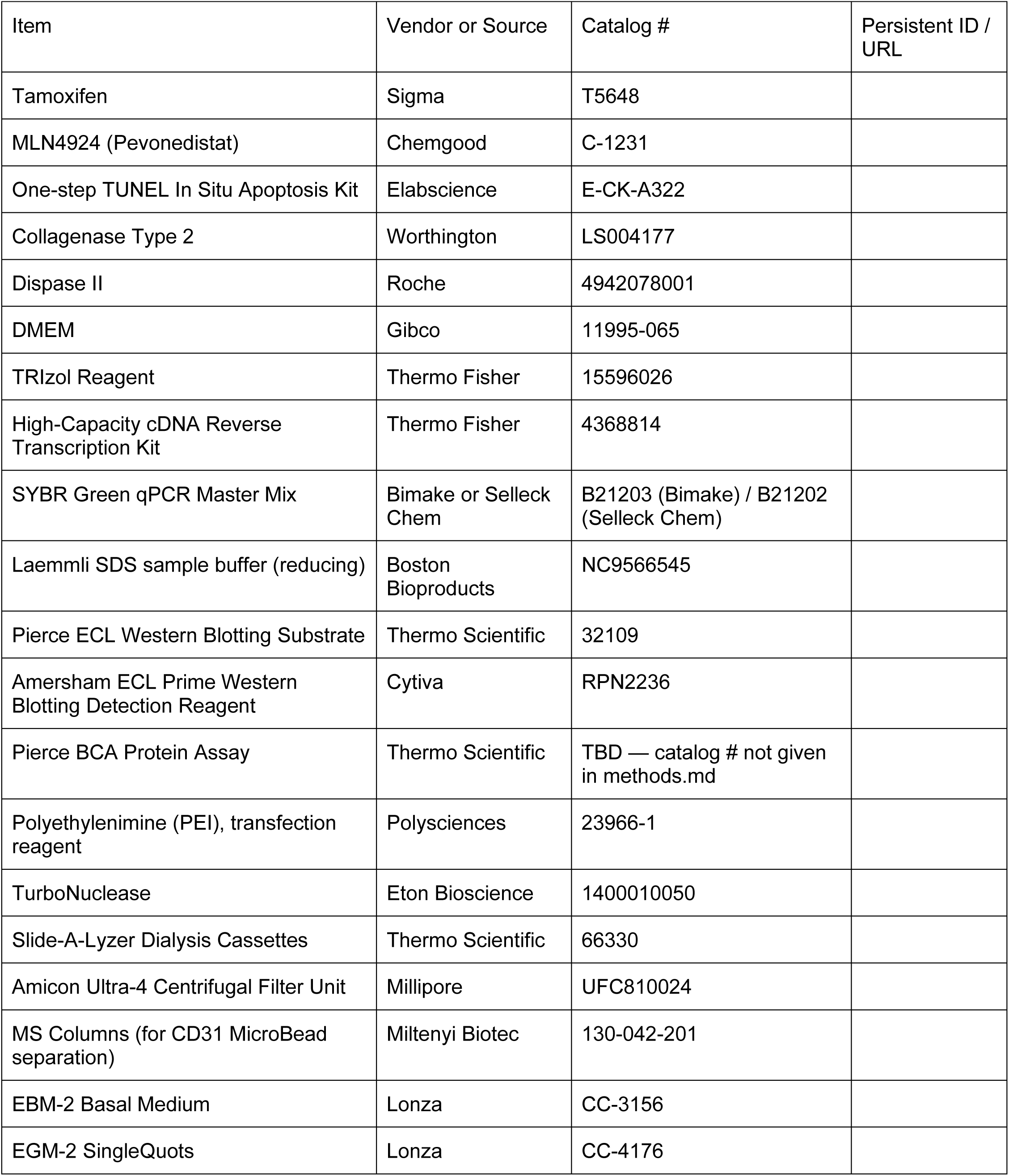

